# Dual role of BdMUTE during stomatal development in the model grass *Brachypodium distachyon*

**DOI:** 10.1101/2024.05.01.592049

**Authors:** Roxane P. Spiegelhalder, Lea S. Berg, Tiago D. G. Nunes, Melanie Dörr, Barbara Jesenofsky, Heike Lindner, Michael T. Raissig

## Abstract

Grasses form morphologically derived, four-celled stomata, where two dumbbell-shaped guard cells (GCs) are flanked by two lateral subsidiary cells (SCs). This innovative form enables rapid opening and closing kinetics and efficient plant-atmosphere gas exchange. The mobile bHLH transcription factor MUTE is required for SC formation in grasses. Yet, if and how MUTE also regulates GC development and if MUTE mobility is required for SC recruitment is unclear. Here, we transgenically impaired BdMUTE mobility from GC to SC precursors in the emerging model grass *Brachypodium distachyon*. Our data indicates that reduced BdMUTE mobility severely affected the spatiotemporal coordination of GC and SC development. Furthermore, while BdMUTE has a cell-autonomous role in GC division orientation, complete dumbbell morphogenesis of GCs required SC recruitment. Finally, leaf-level gas exchange measurements showed that dosage-dependent complementation of the four-celled grass morphology was mirrored in a gradual physiological complementation of stomatal kinetics. Together, our work revealed a dual role of grass MUTE in regulating GC division orientation and SC recruitment, which in turn was required for GC morphogenesis and the rapid kinetics of grass stomata.

## INTRODUCTION

Stomata - microscopic pores at the leaf surface - first appeared with the colonization of dry land and enabled plants to control carbon dioxide (CO_2_) uptake and water vapor loss in leaves sealed with a waxy cuticle (Berry et al., 2010; Chang et al., 2023; Clark et al., 2022). Fossil records suggest that the ancestral stomatal form consisted of two kidney shaped guard cells (GCs) flanking a pore (Edwards et al., 1998). While cellular arrangements of stomata are diverse among the plant kingdom (Gray et al., 2020; Nguyen and Blatt, 2024; Nunes et al., 2020; Rudall et al., 2013), GC morphology in most phylogenetic clades did not derive from its ancestral, kidney-shaped form. This is with the exception of grass GCs that show a unique dumbbell shape with elongated, narrow middle parts (i.e. the central rod) and large bulbous apices (Galatis and Apostolakos, 2004; Spiegelhalder and Raissig, 2021; Stebbins and Shah, 1960). Grass stomata also possess lateral, paracytic (i.e. parallel) subsidiary cells (SCs) (Gray et al., 2020; Nguyen and Blatt, 2024; Nunes et al., 2020; Rudall et al., 2017). Grass SCs are of perigenous origin, meaning they stem from a distinct precursor cell than the GCs.

The dumbbell morphology was suggested to provide a geometric advantage by decreasing the volume to surface ratio and elongating the central pore (Durney et al., 2023; Franks and Farquhar, 2007; Nunes et al., 2020). SCs are suggested to provide molecular and mechanical support to GCs contributing to faster stomatal movements and improved water use efficiency (WUE) (Durney et al., 2023; Franks and Farquhar, 2007; Nunes et al., 2020; Raissig et al., 2017; Raschke and Fellows, 1971; Stebbins and Shah, 1960).

In grasses, epidermal development is organized in linear, semi-clonal cell files, some of which acquire capacity for the formation of stomata (Nunes et al., 2020; Raissig et al., 2016; Stebbins and Shah, 1960). Grass stomatal development starts with a transverse asymmetric division that specifies the young guard mother cell (GMC), which then elongates and induces subsidiary mother cell (SMC) fate in the flanking cell files (Fig. 1A). SMCs then undergo an asymmetric, longitudinal division to form the flanking SCs (Fig. 1A). After SC formation, GMCs divide once symmetrically and longitudinally to form the GC pair before complex morphogenetic processes form the pore, establish cell wall anisotropy and generate the dumbbell morphology (Fig. 1A) (Spiegelhalder and Raissig, 2021). Much like in the well-established plant model *Arabidopsis thaliana* (Lee and Bergmann, 2019; McKown and Bergmann, 2020), stomatal development is guided by the conserved set of key bHLH transcription factors SPEECHLESS (SPCH) (Raissig et al., 2016), MUTE (Raissig et al., 2017; Wang et al., 2019; Wu et al., 2019) and FAMA (Liu et al., 2009; McKown et al., 2023) and their heterodimerization partners INDUCER OF CBF EXPRESSION1 (ICE1) and SCREAM2 (SCRM2) (Raissig et al., 2016).

**Figure 1:**
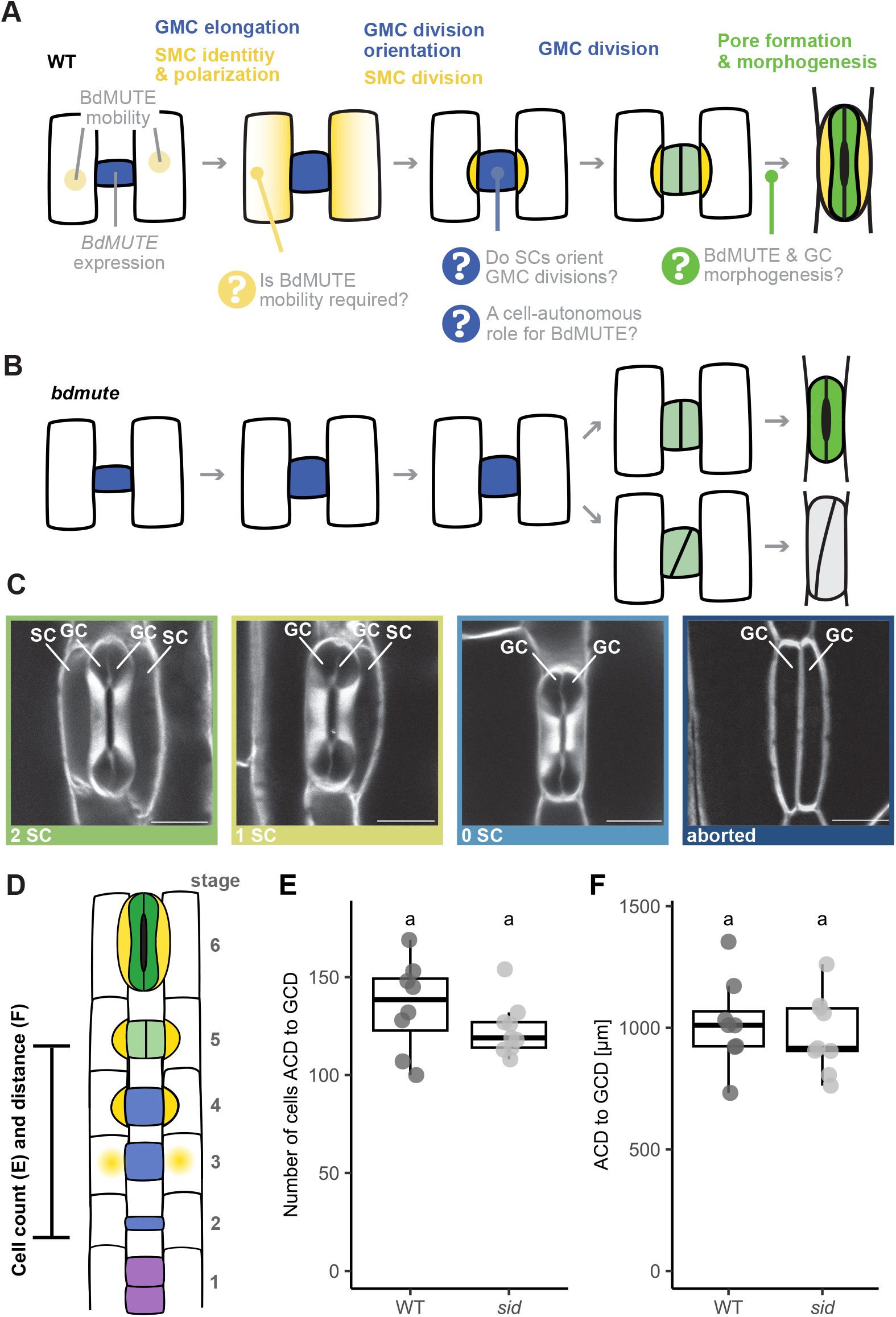
Multiple roles for *BdMUTE* in subsidiary cell (SC) and guard cell (GC) formation in the grass stomatal lineage? (**A**) Schematic displaying stomatal development in wild type (WT). Stomatal development starts with guard mother cell (GMC; blue) formation. *BdMUTE* is expressed in the GMC (light blue nucleus) and moves into the lateral cells (light yellow nucleus) to establish subsidiary mother cells (SMCs). SMCs then polarize (yellow gradient) and divide asymmetrically to form lateral SCs (yellow). Finally, the GMC divides symmetrically to generate two GCs (light green), which form a pore and mature into the characteristic dumbbell shape (dark green). Open questions regarding BdMUTE’s roles and mechanisms are indicated. (**B**) In *bdmute* however, no SMC is established, and no SC division occurs. GMCs, however, mostly divide normally and the GC pair forms a pore and matures into functional complexes (upper GCs, dark green). However, approximately ¼ of the GMCs fail to position the division plane correctly and produce aborted complexes with oblique divisions (lower GCs, gray). (**C**) Different stomatal morphologies found in WT (2 SC), and in *sid*/*bdmute-1* (1 SC, 0 SC and oblique division/arrested phenotype). Labels indicate GCs and SCs. Scale bars = 10μm. Midplane confocal images are of fixed, cleared and Direct Red 23-stained mature stomata. (**D**) Schematic representation of the zone measured in (E) and (F); from first asymmetric cell division (blue, stage 2) to symmetric GC division (light green, stage 5). Developmental stages are indicated; (1) stomatal file initiation, purple, (2) asymmetric division and GMC formation, blue, (3) SMC establishment, yellow gradient, (4) SMC division, yellow, (5) GMC division, light green and (6) GC maturation, dark green. **(E, F)** Number of cells (E) and distance (F) between the first asymmetric cell division (ACD, stage 2) and the first GMC division (GCD, stage 5) in WT and *sid*/*bdmute-1*. n = 5-7 individuals per genotype, 1 or 2 stomatal rows per individual (dots are stomatal rows). Significant differences are indicated with differing letters. Statistical test: Two-sided Student’s t-test (significant if p < 0.05).

The bHLH transcription factor MUTE holds a key role in guiding grass stomatal development (Nunes et al., 2020; Serna, 2020). MUTE’s primary role in grasses is establishing the SC lineage and driving its longitudinal asymmetric division (Raissig et al., 2017; Wang et al., 2019). In the wild model grass *B. distachyon* (Raissig and Woods 2022) and the domesticated cereal grasses maize and rice, MUTE is expressed in GMCs only, but the protein then moves laterally into the neighboring cells thereby establishing SC lineage identity (Raissig et al., 2017; Wang et al., 2019). This would be an elegant yet simple mechanism to establish stomatal identity in non-stomatal cell files. In *B. distachyon*, approximately 75% of *bdmute* stomata form functional, two-celled GC complexes that lack SCs (Raissig et al., 2017). In domesticated grasses like maize and rice, however, not only SC but also GC development completely aborts in *mute* mutants as all of the GMCs fail to divide in the proper orientation and no functional stomata are formed (Wang et al., 2019; Wu et al., 2019). This might be an example for domestication-induced loss of genetic diversity causing a lack of developmental compensation of GC formation. However, much like in domesticated grasses, ∼25% of *bdmute* GMCs in *B. distachyon* show oblique, misoriented division planes, which lead to stomatal abortion (Raissig et al., 2017). And finally, even though functional two-celled complexes form in *B. distachyon*, the GCs are shorter and develop less of the dumbbell-shaped morphology compared to their wild-type stomata flanked by SCs.

This raises many open questions regarding the role and functionality of grass MUTE (Fig. 1A). First, is there a cell-autonomous role for MUTE in guiding GC development as suggested by the partial or full GC arrest in wild and domesticated grasses? Is this role primarily pre-mitotic by guiding division plane orientation or also post-mitotic by driving GC differentiation? Or are the observed division plane orientation defects a mere consequence of the lack of SC recruitment, which could provide positional information and reinforce GMC division planes? Second, is MUTE mobility indeed required for MUTE function? Or are downstream mobile factors or cell-to-cell signaling responsible for SC formation? Third, is the impaired morphogenesis of *bdmute* GCs a consequence of missing SCs or a missing, cell-autonomous role of BdMUTE in the GC lineage?

To address these open questions, we rescued *sid/bdmute-1* with a 3xGFP-MUTE construct with impaired mobility (*BdMUTEp:3xGFP-BdMUTE*, M3GM). Scoring the 3xGFP-BdMUTE’s ability to rescue the GC and SC defects, we were able to show that BdMUTE acts cell-autonomously in guiding the GMC division, independent of its role in SC formation. GC morphogenesis, however, required the presence of SCs, showing that both cell- and non-cell-autonomous aspects influence GC development in *B. distachyon*. Furthermore, our data suggested that physical presence of MUTE in SMCs (i.e. MUTE mobility) is relevant for SC recruitment. 3xGFP tags merely reduced but did not abolish BdMUTE mobility. Therefore, the strongest 3xGFP-BdMUTE line was able to fully rescue both GC and SC defects in mature leaves, but caused a severe developmental delay of SC recruitment and spatiotemporal disconnect between GC and SC divisions. Together, our data suggest a dual role of *BdMUTE* to spatiotemporally synchronize the development of the two stomatal cell types to build a physiologically superior, four-celled stomatal complex in grasses.

## MATERIAL AND METHODS

### Plant material and growth conditions

Bd21-3 *Brachypodium distachyon* was used as wild type (WT). 2^nd^ leaf confocal microscopy was conducted on plate-germinated seedlings that were sterilized with 20% bleach, 0.1% Triton X-100, washed and placed on ½ MS Murashige & Skoog (Duchefa) plates. After 2 days of vernalization and stratification (in dark, 4°C), these seeds were placed into a 28°C chamber with 16 hr light: 8 hr dark cycle, 110 μmol photosynthetic active radiation (PAR) m^−2^ s^−1^ for 5-6 days until imaging.

After 2^nd^ leaf microscopy, the seedlings were planted in soil, consisting of four parts ED CL73 (Einheitserde) and one part Vermiculite and grown in a greenhouse or growth chamber with 18 hr light:6 hr dark cycle (250–350 μmol PAR m^−2^ s^−1^; day temperature = 28°C, night temperature = 22°C); (also described in Haas and Raissig, 2020; Nunes et al., 2022). The *sid/ bdmute-1* line described in (Raissig et al., 2017) was used as a mutant line and background for reporter constructs.

### Generation of reporter constructs

Reporter constructs were used as described in (Raissig et al., 2017) (*BdMUTEp:YFP-BdMUTE, BdMUTEp:Bd-MUTE-YFP*) or generated using the GreenGate cloning system (Lampropoulos et al., 2013). For Greengate cloning, *BdMUTE* genomic region (BdiBd21-3.1G0240400) was amplified from Bd21-3 WT genomic DNA with priT-N7+priTN8. The *ZmUBI* promoter and *BdMUTE* promoter were generated and used as described in (Nunes et al., 2023). A backbone-specific GreenGate overhang was created in the process and the product was used to generate the respective GreenGate entry vector.

The entry modules, pGGD002 (D-dummy) and pGGE001 (*rbcS* terminator) are described in (Lampropoulos et al., 2013) and pGGZ004 described in (Lupanga et al., 2020). The entry module pGGB_3xGFP was generously provided by Jan Lohmann (Centre for Organismal Studies, Heidelberg).

The expression vectors *ZmUBIp:3xGFP-BdMUTE* and *BdMUTEp:3xGFP-BdMUTE* were assembled using the Green-Gate cloning system. The 6 entry modules (pGGA_specific promoter (*MUTEp* or *ZmUBIp*); pGGB_3xGFP; pGGC_BdMUTE_genomic; pGGD_dummy; pGGE_terminator and pGGF_resistance (Zhang et al. 2022) were repeatedly digested and ligated with the destination vector pGGZ004 during 50 cycles (5 min 37°C followed by 5 min 16°C) followed by 5 min at 50°C and 5 min at 80°C for heat inactivation of the enzymes. All final constructs were test digested and the generated GreenGate overhangs were Sanger sequenced.

### Generation of transgenic lines

Generation of transgenic lines was performed as described in (Zhang et al., 2022). In brief, embryonic calli were cultivated from isolated embryos from Bd21-3 and *sid/bdmute-1* on callus induction media (CIM; per L: 4.43 g Linsmaier & Skoog media (LS; Duchefa), 30 g sucrose, 600 μl CuSO4 (1 mg/ml, Sigma/Merck), 500 μl 2,4-D (5 mg/ml in 1M KOH, Sigma/Merck), pH 5.8, plus 2.3 g of Phytagel (Sigma/Merck)) for 3, 2 and one weeks, interspaced with transferral and splitting to fresh CIM media. Cultivation of the forming calli took place at 28°C, in the dark.

For transformation of the calli, AGL1 *A. tumenfaciens* with the desired construct were scraped off selection plates and dissolved in liquid CIM media (same media as above without the phytagel) with freshly added 2,4-D (2.5 mg/ml final concentration), Acetosyringone (200 mM final concentration, Sigma/Merck), and Synperonic PE/F68 (0.1% final concentration, Sigma/Merck). Around 100 calli (approximately two plates) were incubated in AGL1 in solution with OD600 = 0.6 for 15 minutes.

Afterwards, the calli were dried off on sterile filter paper, incubated three days at room temperature in the dark and then moved to selection media (CIM + Hygromycin (40 mg/ml final concentration, Roche) + Timentin (200 mg/ml final concentration, Ticarcillin 2NA & Clavulanate Potassium, Duchefa)). On selection media, the calli were incubated for one week and then moved to fresh selection plates and incubated for two more weeks at 28°C in the dark. Next, the calli were moved to callus regeneration media (CRM; per L: 4.43 g of LS, 30 g maltose (Sigma/Merck), 600 μl CuSO4 (1 mg/ml), pH 5.8, plus 2.3 g of Phytagel, Timentin (200 mg/ml final concentration), Hygromycin (40 mg/ml final concentration), and sterile Kinetin solution (0.2 mg/ml final concentration, Sigma/Merck)). The transformed calli were incubated at 28°C in a 16h light: 8h dark cycle (70-80 μmol PAR m^-2^ s^-1^). Shoots > 1cm that formed after 2-6 weeks in the light on regenerating calli were transferred to rooting cups (Duchefa) containing rooting media (per L: 4.3 g Murashige & Skoog including vitamins (Duchefa), 30 g sucrose, pH 5.8, 2.3 g Phytagel, Timentin 200 mg/ml final concentration) and grown at 28°C in a 16h light: 8h dark cycle (70-80 μmol PAR m^-2^ s^-1^). Once roots had formed, plants were moved to soil (4 parts ED CL73 (Einheitserde), 1 part Vermiculite) and grown in a greenhouse/growth chamber with 18 h light: 6 h dark cycle (250-350 μmol PAR m^-2^ s^-1^).

### Microscopic analysis of SC and GC complementation phenotypes

3^rd^ leaves for quantitative assessment of rescue phenotype and for apex/middle ratio quantifications were collected from 3 weeks old, soil-grown plants. For adult leaf phenotyping, the leaf used for LI-6800 gas exchange measurements was collected. The leaves were fixed in 7:1 ethanol:acetic acid. Before imaging, leaf tissue was rinsed twice in water and mounted on slides in Hoyer’s solution (Sharma, 2017). The abaxial side was imaged using a 40x objective on a Leica DM5000B or Leica DM2000 microscope with differential interference contrast (DIC) imaging (Leica Microsystems).

3D-stacks of mature stomatal complexes were imaged on mature leaves of plants older than 5 weeks. Fixing and staining was adapted after (Ursache et al., 2018) which in turn is based on (Kurihara et al., 2015). The leaf tissue was fixed with 4% PFA (paraformaldehyde) in 1 x PBS by pressure-injection with a needleless syringe and following 1 h incubation in 4% PFA with gentle agitation. The leaf was subsequently washed twice with 1x PBS and transferred to ClearSee solution (Xylitol [final concentration 10% (w/v)] (Sigma), sodium deoxycholate [final concentration 15% (w/v)] (Sigma), Urea [final concentration 25% (w/v)] (Sigma) in H_2_O). The leaf tissue was cleared in ClearSee at least 24-72 h with gentle agitation and stained with 0.1% Direct Red 23 (Sigma) in ClearSee overnight. To remove excess staining the leaves were washed in ClearSee for at least 1 h before imaging.

Imaging of 3D mature leaves was carried out at a Leica SP8 confocal microscope with 20% Argon laser and 561 nm excitation and detection at 580-615 nm at an intensity that yielded slight oversaturation for better assessment of the 3D structure. Gain was corrected in z-stacks using the z-compensation tool to acquire clear images in all z-planes. Stacks were imaged at resolution of 1024x1024 pixels, in 0.1 μm z-steps with a line average of 2.

### Microscopic analysis of reporter lines

For confocal imaging of reporter constructs, the developmental zone (first 2 mm) of the 2^nd^ leaf of 5-7 days post germination, plate-germinated seedlings were used. Samples were stained with propidium iodide (PI, 10 μg/ml, Thermofisher) for approximately 5 min to visualize cell walls, mounted in H_2_O on a slide and imaged with a 63x glycerol immersion objective on a Leica TCS SP8 microscope or a Leica Stellaris 5 microscope (Leica Microsystems). On the SP8 we imaged using an Argon laser at 20% power, at the Stellaris 5 a white light LED at 85% laser power was used. At both microscopes, two high-sensitivity spectral detectors (HyD) were used to detect fluorescence emission (Leica Microsystems).

Laser intensity for fluorophore excitation was set to allow clear PI and fluorophore signals. Laser intensity was kept constant for quantification of the intensity, excitation of GFP with 488 nm at 15% intensity and 150% gain and PI excitation at 514 nm, adjusted for cell wall visibility. These settings allowed to still visualize the lowest expressing 3xGFP line but led to saturation of the highest expressing line. The two YFP lines were excited with 12% intensity, 100% gain and plotted separately, as different fluorophores are not comparable. For all confocal microscopy, midplane images were used except when specified otherwise.

CTCF (corrected total cell fluorescence, El-Sharkawey, 2016) measurements were performed on confocal images of 2^nd^ leaf developmental zones. Images were acquired in stacks of 0.5 μm step size, 1024x1024 pixels and 2x line average. The average intensity of 3 adjoining slices with highest intensity of those stacks were taken to allow correction for nuclear position. Then the nucleus was traced with the Fiji polygon selection tool and the integrated density of the signal measured. 1-2 background measurements were taken close to, but outside of, the nuclear signal. CTCF was calculated as: integrated density – (area of selected cell X mean fluorescence of background readings).

For mobility ratio measurements, images of M3GM++ and MYM were acquired at the Stellaris 5 confocal microscope with settings that allowed a visualization of SMC nuclear signal and GMC cytoplasmic signal. Those were 489 nm laser with 37.4% intensity and 200% signal gain for the 3xGFP line and 515 nm laser with 3.56% intensity and 75% signal gain for the YFP line. Additionally, for the GFP line, the PI signal was excited with the 549 nm laser. When imaging YFP lines, PI was also excited with the 515 nm laser used to excite YFP.

Images were acquired in stacks of 4 slices with 0.33 μm step size and 1024x1024 pixels.

The average intensity of 2-3 slices of those stacks were taken to allow correction for nuclear position. Then a constant area was selected for each stomatal complex and the integrated density was measured for the SMC_(nuclear)_ and GMC_(cytoplasmic)_ signal. As a proxy for protein mobility, the ratio of SMC_(nuclear)_ and GMC_(cytoplasmic)_ signal was used. This also normalized for the difference in laser intensities.

Staging of the SC recruitment was performed on MYM and M3GM++ 2^nd^ leaf developmental zones. For staging relative to the 1^st^ GC division, up to 15 stomatal complexes apically and basally of the 1^st^ division in one stomatal row were analyzed regarding their GC division status (divided yes or no) and their number of recruited SCs (0/1/2). One stomatal cell file per individual was analyzed in a total of 10 individuals.

SC recruitment linked to the length/width ratio of the GMC/GC was also performed on MYM and M3GM++ 2^nd^ leaf developmental zones; here the length and width of the undivided or divided GMC was measured with the Fiji line tracing tool (perpendicular to the cell wall) and the complexes were analyzed regarding their GC division status (divided yes/no) and their number of recruited SCs (0/1/2). 129-191 stomatal complexes in 2-6 individuals were analyzed.

### Quantification of stomatal morphology

For quantification of the SC and GC division complementation phenotypes in WT, MYM, M3GM (++, +, - and -), MMY and *sid/bdmute-1* 19-44 images of 3^rd^ leaves were acquired with the 40x objective at the DIC microscope. For 5-12 individuals the stomatal phenotypes were counted in the four categories “2 SC, 1 SC, 0 SC, aborted”. Per individual, 74-244 stomatal complexes were counted.

GC morphology was quantified by measuring stomatal complexes of the M3GM-line flanked by zero, one or two SCs. The length of the GCs was measured along the longitudinal middle cell wall. Transverse GC width in the apices was measured at the widest and in the narrow central rod at the middle at the narrowest point. All measurements were performed with the line tracing tool in Fiji.

To correct for focal plane bias, 3D stacks of mature, Direct Red 23-stained stomatal complexes flanked by zero, one or two SCs of the M3GM-line were acquired. Of those complexes, XZ optical cross sections for each GC at three points of the Y axis and one YZ optical section of each GC were taken. The XZ optical cross sections were made at the widest points of the apical and basal bulbous apices and the narrowest point of the middle rods. The area of the GC at these three sections was measured using the polygon selection tool in Fiji and the ratio of average apex area to middle area was taken as a proxy for GC dumbbell shape.

Quantifications of cell number and total tissue length between the first stage 2 cell (transverse asymmetric division) and the first stage 5 cells (longitudinal symmetric GMC division) comparing WT and *sid/bdmute-1* were performed in developmental zones of plate-grown 2^nd^ leaves. 5-7 leaf zones per genotype were stained with PI and imaged using a 63x glycerol immersion objective at a SP8 confocal microscope (Leica Microsystems). Images were captured along the developmental zone of each sample starting at the first asymmetric division in the stomatal file and moving up to the first symmetric division of the GC precursors. Individual images along the stomatal rows were taken with at least 20% overlap and potential landmarks to later facilitate stitching and aligning. Images were processed with Fiji (Schindelin et al., 2012). Images were stitched pairwise using the “stitching” plugin and manually whenever pairwise stitching was not possible. To quantify the cell number between developmental zones, cells were counted using the multi-point tool and the distances between first ACD and GCD calculated.

Division orientation of GMCs in *sid/bdmute-1*, MMY, M3GM- - and WT was assessed using confocal imaging. Images were obtained of PI-stained developmental zones from 2^nd^ leaves of plate-germinated seedlings using the 63x objective. Both the imaging and the analysis was done fully anonymized (i.e. neither the person doing the microscopy nor the person doing the analysis knew the genotype of the sample). A straight line was drawn connecting the two apical or the two basal lateral corners of the divided GMC of the stomatal complex. Then, the left and right distance were measured as distance from the corner to the intersecting GMC division plane on the previously drawn line (Fig. 4B). For each stomatal complex, the smaller distance was divided by the bigger distance to obtain the ratio of the division plane skewness.

All image analysis and processing was carried out in Fiji (Schindelin et al., 2012). Images shown in Figs 1, 3, 4 and 5 were processed with background reduction (rolling ball 50) and brightness and contrast adjustments. For pictures shown in Fig. 2 only the PI channel was adjusted. No post-processing was performed on images that were used for quantification.

**Figure 2:**
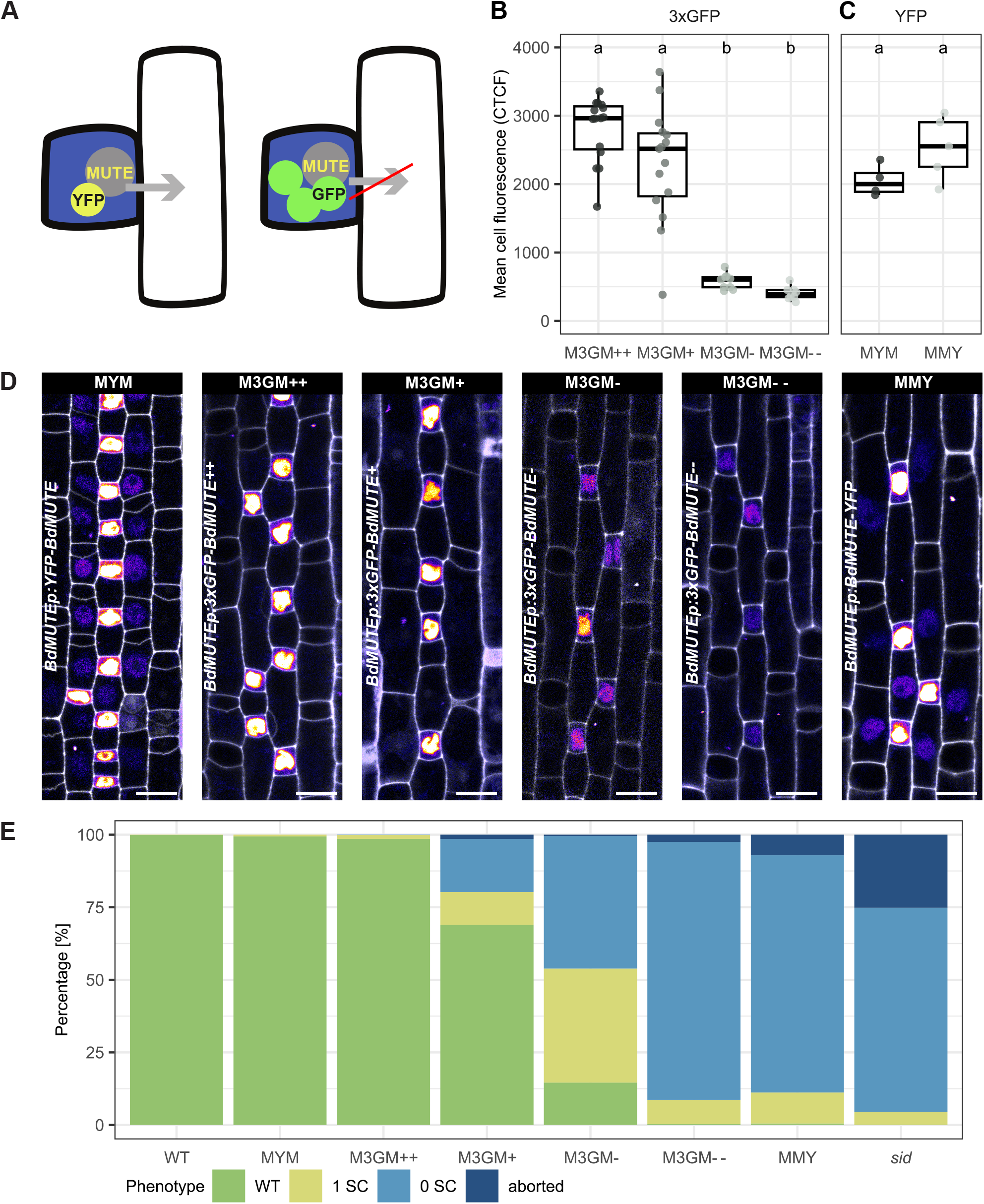
Expression of mobility impaired 3xGFP-BdMUTE rescues *sid*/*bdmute-1* in a dose-dependent manner. (**A**) Schematic displaying the reporter constructs. Single tagged BdMUTE (*sid/bdmute-1*;*BdMUTEp:YFP-BdMUTE*, MYM, yellow+gray) can move from GMCs (blue) into lateral cell files (white). A triple GFP tag (*sid/bdmute-1*;*BdMUTEp:3xGFP-BdMUTE*, green+gray) impairs BdMUTE mobility and should restrain BdMUTE to GMCs (blue). (**B**) GFP fluorescence measured in *BdMUTEp:3xGFP-BdMUTE* (M3GM) reporter lines in the *sid*/*bdmute-1* background. Corrected total cell fluorescence (CTCF) of the average of 3 slices of a z-stack around the midplane of the GMC was measured; all images taken with constant laser settings. n = 8-15 individuals per genotype and 49-105 stomata per genotype (dots are individuals). Significant differences are indicated with differing letters. Statistical test: ANOVA followed by Tukey’s HSD test (alpha = 0.05). (**C**) YFP fluorescence of *BdMUTEp:YFP-BdMUTE* (MYM) and *BdMUTEp:BdMUTE-YFP* (MMY) in the *sid*/*bdmute-1* background. CTCF of the average of 3 slices of a z-stack around the midplane of the GMC was measured; all images taken with constant laser settings. n = 4-5 individuals per genotype and 29-31 stomata per genotype (dots are individuals). Significant differences are indicated with differing letters. Statistical test: ANOVA followed by Tukey’s HSD test (alpha = 0.05). (**D**) Representative midplane confocal images of stage 3 leaf zones of the lines shown in (B) and (C) with signal intensity of reporter proteins shown as “fire” heatmap and propidium iodide (PI)-stained cell walls shown in gray. MUTE in GMCs shows different intensities in the four GFP lines, mobile signal in the lateral cell files is only visible in the single-tag YFP lines. GFP was imaged with 15% laser and 150% gain, MYM with 12% laser and 100% gain, MMY with 10% laser and 150% gain. Scale bars = 10 μm. (**E**) Stomatal phenotypes in the mature 3^rd^ leaf of WT, *sid/bdmute-1* and *sid/bdmute-1* complemented with the different reporter lines shown in (D). n = 5-12 individuals per genotype and 864-1450 stomata per genotype. Same data can be found as boxplots in Fig. S2.

**Figure 3:**
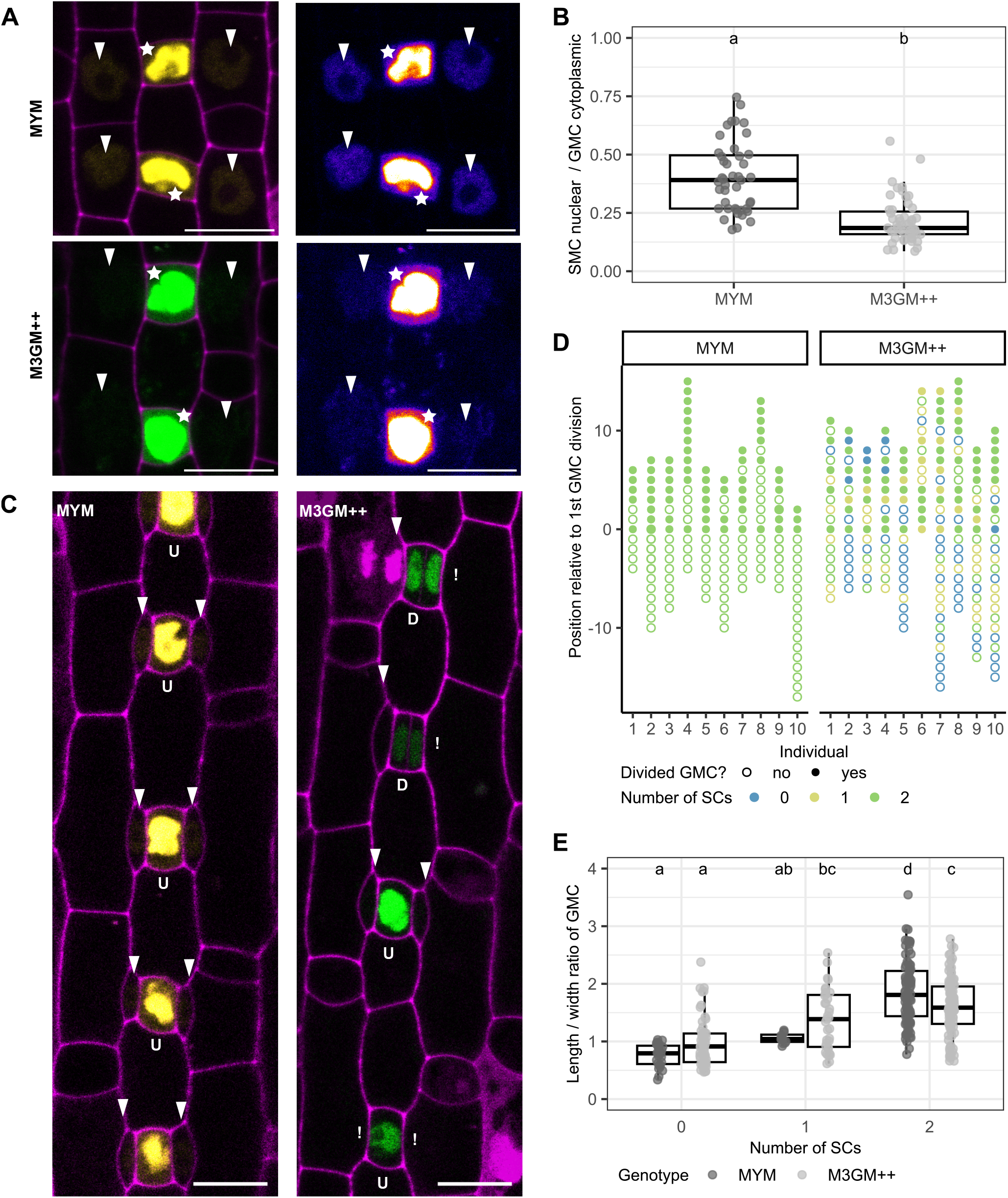
A triple GFP tag reduces BdMUTE mobility leading to a temporal delay in SC recruitment. (**A**) Representative images of guard mother cells (GMCs; stars) and lateral subsidiary mother cells (SMCs; arrowheads) of *sid/bdmute-1*;*BdMUTEp:YFP-BdMUTE* (MYM; top panels) and *sid/bdmute-1*;*BdMUTEp:3xGFP-BdMUTE* (M3GM++, bottom panels). Cells are imaged with laser intensities that assured non-saturated intensity in cytoplasm of GMCs and visible signal in SMC nuclei. YFP/GFP in yellow/green and propidium iodide (PI)-stained cell walls in purple in the left panels and heatmap projection of YFP/GFP intensity in right panels; Scale bars = 10 μm. (**B**) Ratio of integrated density of average SMC nuclei signal and average GMC cytoplasmic signal. n = 2-3 individuals per genotype and 48-58 stomatal complexes per genotype (dots are stomata). Significant differences are indicated with differing letters. Statistical test: ANOVA followed by Tukey’s HSD test (alpha = 0.05). (**C**) Midplane confocal images of 2^nd^ leaf developmental zones in MYM and M3GM++; undivided (“U”) and divided (“D”) GMCs and present SCs (arrowhead) or missing SCs (!) are indicated. Cell walls are stained with PI (magenta), fluorophores shown in yellow (YFP) and green (3xGFP). Scale bars = 10 μm. (**D**) Division and SC recruitment status of GMCs apically and basally relative to the first GMC division in M3GM++ and MYM. Each dot represents one cell and is aligned according to its position relative to the first GMC division; filled dots indicate divided GMCs, empty dots indicate non-divided GMCs; color represents the number of SCs recruited (green = 2 SCs; yellow = 1 SC; blue = no SCs). Each column represents a single stomatal row (n = 10 individuals per genotype). (**E**) Length/width ratio of GMCs in 2^nd^ leaf developmental zones in MYM and M3GM++ flanked by zero, one and two SCs, respectively. Color indicates genotype; n = 2-6 individuals per genotype and 129-191 stomata per genotype. Significant differences are indicated with differing letters. Statistical test: ANOVA followed by Tukey’s HSD test (alpha = 0.05).

**Figure 4:**
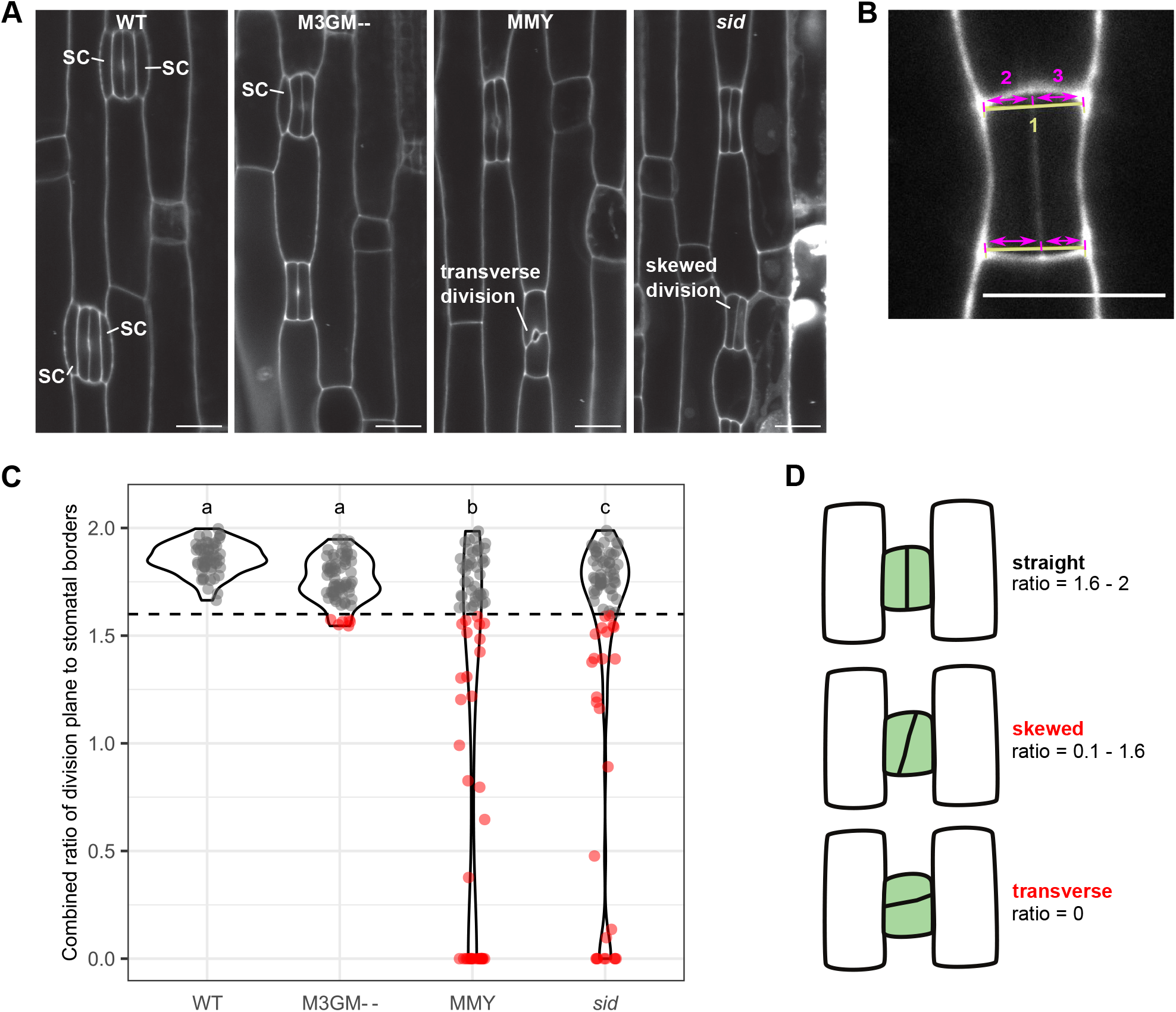
Low BdMUTE levels in guard mother cells (GMCs) mostly rescue the GMC division plane orientation defects. (**A**) Representative images of recently divided GMCs in wild type (WT), *sid/bdmute-1*;*BdMUTEp:3xGFP-BdMUTE* (M3GM - -), *sid/bdmute-1*;*BdMUTEp:BdMUTE-YFP* (MMY), and *sid*/*bdmute-1*. Midplane confocal images of developmental zones, stained with propidium iodide (PI). Scale bars = 10μm. (**B**) Schematic of division plane orientation measurements. A straight line was drawn to connect the outward corners of the GMC at the apical and basal side, respectively (1). The distance from each corner to the intersection with the GMC division plane was measured along the previously drawn line (2 and 3). The smaller distance was divided by the longer distance to obtain the “skewness” ratio of the GMC division plane for the apical (Fig. S3A) and the basal side (Fig. S3B). **(C)** Total “skewness” ratios per GMC as a sum of apical and basal “skewness” ratios. A total ratio < 1.6 (dashed horizontal line) represents a skewed division (indicated as red dots). n = 7-8 individuals per genotype and 66-82 stomata per genotype (dots are stomata). Significant differences are indicated with differing letters. Statistical test: ANOVA followed by Tukey’s HSD test (alpha = 0.05). **(D)** Schematic representation of the division phenotypes depicted in (C). “Straight” division represented by ratios > 1.6 (i.e. all WT stomata) were used as a WT cutoff. GCs with a division plane orientation < 1.6 and > 0 were classified as “skewed” and a transverse division is shown as ratio = 0.

**Figure 5:**
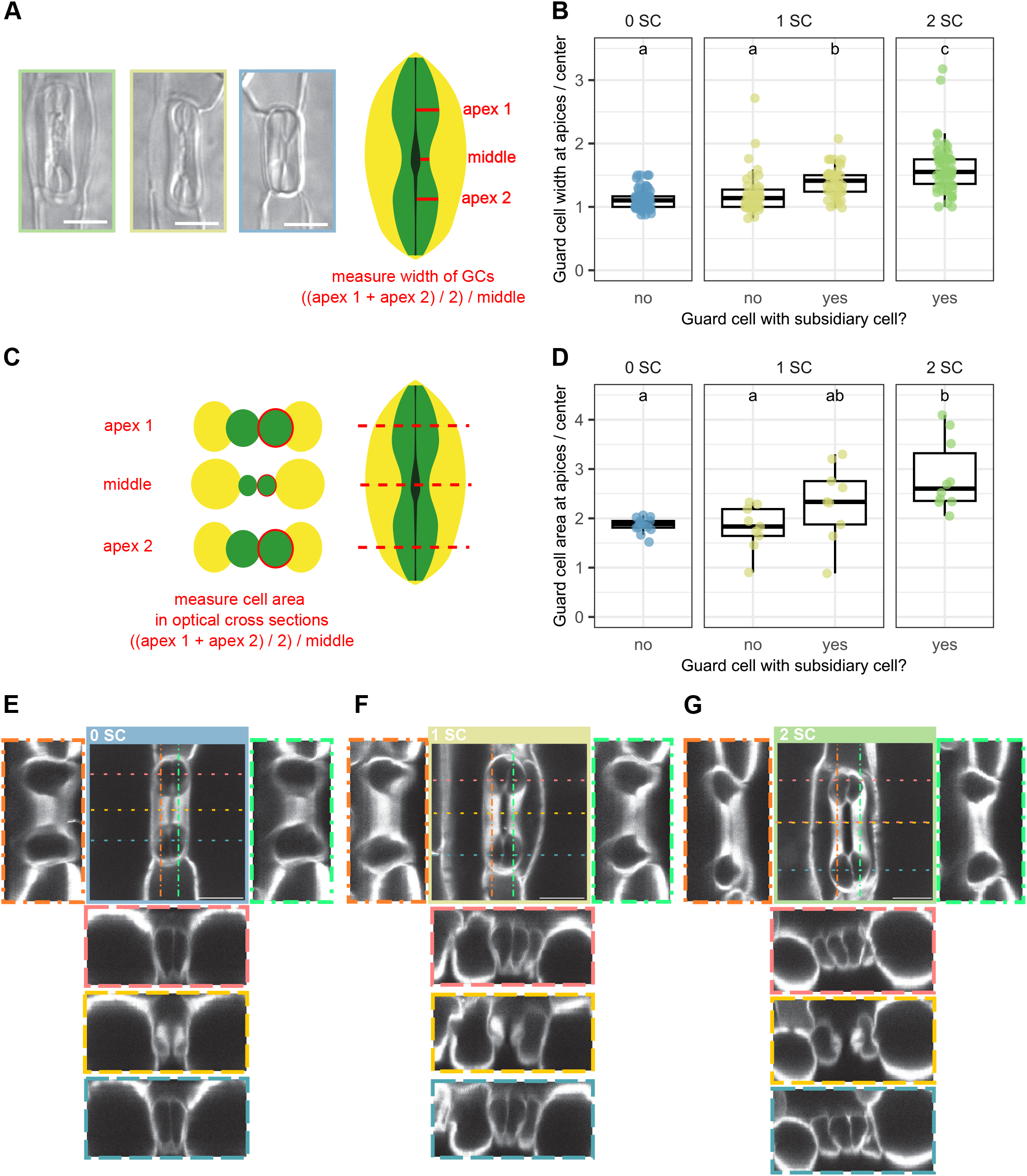
Presence of subsidiary cells (SCs) influences guard cell (GC) morphology. (**A**) Differential interference contrast (DIC) images of different stomatal morphologies of the M3GM-line, showing complexes with two, one or no SCs (green, yellow and blue box, respectively). A schematic indicates how these complexes were measured to calculate a curvature ratio in (B). Scale bars = 10 μm. (**B**) GC width at the apices / middle ratio of M3GM-GCs flanked by either zero, one or two SCs. Ratio indicates curvature of GCs; each data point represents one cell, measurements performed on DIC images as shown in (A). n = 2 individuals and 16-36 stomata per phenotype. Significant differences are indicated with differing letters. Statistical test: ANOVA followed by Tukey’s HSD test (alpha = 0.05). (**C**) Schematic illustrating how optical cross sections taken from 3D confocal stacks (shown in E-G) were analyzed to calculate curvature ratio of cross section areas in (D). (**D**) Quantification of average apex / middle area ratio of GCs with zero, one or two SCs in M3GM-. Measured were XZ cross sections as shown in (C) of 3D stacks of confocal images (see E-G). Each dot represents one GC. n = 21 stomatal complexes. Significant differences are indicated with differing letters. Statistical test: ANOVA followed by Tukey’s HSD test (alpha = 0.05). (**E-G**) 3D confocal stacks of mature stomatal complexes of the M3GM-line without SCs (E), with one SC (F) and with two SCs (G). XZ Cross sections through the apices and middle and YZ sections are displayed and indicated by dashed, colored lines and respective frames. Scale bars = 10 μm. Confocal images are of fixed, cleared and Direct Red 23-stained stomata.

### Quantification of stomatal physiological responses

The LI-6800 Portable Photosynthesis System (LI-COR Biosciences Inc.) infra-red gas analyzer was used to obtain leaf level gas exchange measurements. As described in (Nunes et al., 2022), the youngest, fully expanded leaves of 3 to 4 week old soil-grown plants were measured using the following conditions in the chamber system: flow rate, 500 μmol s^-1^; fan speed, 10’000 rpm; leaf temperature, 28°C; relative humidity (RH), 40%; [CO_2_], 400 μmol mol^-1^. Photosynthetic active radiation (PAR) was set to 1000 - 100 - 1000 - 0 μmol m^-2^ s^-1^, 20 min per step. Every 60 s the system automatically logged gas exchange and environment conditions including absolute stomatal conductance (*g*_sw_) and carbon assimilation (*A*). In total 26 WT individuals due to repeated paired experiments and 4-6 individuals for the M3GM lines (++, +, (*A*) and - -), MYM, MMY and *sid/bdmute-1* were used and the respective leaf section that was inside the Li-6800 chamber was collected in 7:1 ethanol:acetic acid for fixation and clearing for subsequent microscopy (see above). All measurements were corrected in the respective LI-6800 excel output file by individual leaf area calculated as leaf width multiplied by the chamber diameter. The excel files were used as input for the licornetics R package (version 2.1.2, Berg, 2024), available on Github: https://github.com/lbmountain/licornetics) to calculate intrinsic water-use efficiency (iWUE; calculated as *A* divided by *g*_sw_), average the data across individuals, divide the values by stomatal density to obtain values per stoma and visualize it.

To quantify the complementation phenotypes, 9-10 DIC images per individual were taken using the 40x objective. Phenotypes were categorized as “2 SC”, “1 SC”, “0 SC” and “aborted” (see above).

These images were also used to calculate stomatal densities (stomata per mm^2^) and physiological parameters were corrected by mean stomatal density per line in the licornetics package to obtain values per stoma.

### Calculation of stomatal density and stomatal index

Leaves of 10 individuals of WT, M3GM++, MYM and *sid/bdmute-1* were collected 12 and 28 days after germination on soil. Total number of cells and stomatal complex numbers were counted in DIC images of 4 fields of view (20x objective) per individual. Stomatal density was calculated as stomata per mm^2^ and stomatal index as stomatal complexes divided by the total cell number per field of view.

### Statistical analysis and data visualization

Downstream analysis of spreadsheet data files and visualization was conducted in R Studio (version 2023.12.1, RStudio Team and RStudio, PBC, Boston, MA, 2023) using R (version 4.3.1, R Core Team and R Foundation for Statistical Computing, Vienna, Austria, 2023). Statistical analysis was done using either Student’s t-Test or ANOVA (stats package version 4.3.1, R Core Team and R Foundation for Statistical Computing, Vienna, Austria, 2023) and Tukey’s HSD test (agricolae package version 1.3-6, de Mendiburu, 2023). Further packages required for file management, calculations and final visualization were: readxl (version 1.4.3, Wickham and Bryan, 2023), tidyverse (version 2.0.0, Wickham et al., 2019), ggpubr (version 0.6.0, Kassambara, 2023), ggtext (version 0.1.2, Wilke and Wiernik, 2022) and MetBrewer (version 0.2.0, Mills, 2022).

### Data availability

The leaf area-corrected LI-6800 gas exchange (Li-6800-files_Spiegelhalder-et-al_2024.zip), the quantitative data tables used as input for analysis and plotting in R (MUTE2024_ Data.xlsx), and the complete R script used for calculations and visualizations (Spiegelhalder-et-al_2024_BdMUTE.R) are available on Github (https://github.com/lbmountain/Spiegelhalder-et-al_2024_BdMUTE).

## RESULTS

### 3xGFP-BdMUTE rescues *sid/bdmute-1* in a dose-dependent manner

MUTE reporters in all grasses tested showed very strong signal in GMCs and young GCs (i.e. where it is expressed), and much weaker signal in SMCs and young SC (i.e. where it moves to) (Raissig et al., 2017; Wang et al., 2019). In *sid/ bdmute-1*, no SCs are recruited and 25.2% ± 4.4% of GMCs fail to divide properly and abort (Fig. 1B,C, Fig. 2E, Fig. S2). Both phenotypes are fully rescued by complementation with a single mCitrine-tagged BdMUTE under its endogenous promoter (*BdMUTEp:mCitrine-BdMUTE* in *sid/bdmute-1*; Fig. 2E, Fig. S2) (Raissig et al., 2017). Yet, if SC recruitment is needed to time and/or orientate the GMC division, whether BdMUTE protein in SMCs (i.e. BdMUTE mobility) is indeed required to induce SMC divisions, and whether BdMUTE has a GC-autonomous role during GC division and morphogenesis remained to be determined (Fig. 1A).

To test if SC recruitment is required to timely and accurately induce GMC division we quantitatively analyzed the developmental leaf zone of *sid/bdmute-1* (Fig. 1D-F). In both WT and *sid/bdmute-1* leaf zones the number of cells and distance between stage 2 (transverse divisions establishing GMCs) and stage 5 (after GMC division) cells are equal (Fig. 1D-F). This suggested that SC recruitment is not required to time GMC division and indicated at least partial independence between SMC and GMC divisions. Yet, whether BdMUTE presence is indeed required in lateral cell files for SC recruitment and whether SC recruitment or a GC-autonomous role of BdMUTE orients the longitudinal, symmetric GMC divisions remained unknown.

To tackle these questions, we generated a mobility-impaired, but functional BdMUTE construct (*BdMUTEp:3xGFP-BdMUTE*; Fig. 2A) and assessed its ability to rescue different aspects of GC and SC lineage defects in *sid/bdmute-1*. 3xGFP-BdMUTE under its endogenous promoter (*BdMUTEp:3xGFP-BdMUTE*) is supposedly immobile and restricted to the GC lineage, where it is expressed, which should allow observing GMC-autonomous roles of BdMUTE. The use of bulky fluorescent protein tags (i.e. adding three (3xFP) instead of just one (1xFP) fluorescent protein) is an established system to prevent protein mobility (Andersen et al., 2018; Clark et al., 2016; Goretti et al., 2020). To test the functionality of 3xGFP-BdMUTE, we first used an overexpression approach (WT;*ZmUBIp:3xGFP-MUTE*) and observed excessive SC-like divisions (Fig. S1) similar to those observed in single fluorophore overexpression lines (Raissig et al., 2017). We then isolated four independent *BdMUTEp:3xGFP-BdMUTE* (M3GM) lines in the *sid/bdmute-1* background and observed severe expression variability (Fig. 2B-D). We named the lines according to their expression levels: M3GM- - for the weakest expressing line, M3GM- for the weak expressing line, M3GM+ for the strong expressing line, and M3GM++ for the strongest expressing line (Fig. 2B,D). In addition to these four lines, we used wild type (Bd21-3 = WT), *sid/bdmute-1* and *sid/bdmute-1* transformed with the mobile, yet non-functional C-terminally tagged MUTE (*sid/ bdmute-1*;*BdMUTEp:BdMUTE-mCitrine* = MMY) and the mobile, fully functional N-terminally tagged rescue construct (*sid/bdmute-1*;*BdMUTEp:mCitrine-BdMUTE* = MYM) (Raissig et al., 2017) as controls (Fig. 2 C-E). We quantified corrected total cell fluorescence (CTCF) to determine the expression strength of these different lines using the same laser intensities for the GFP lines and a comparable laser setting for YFP lines (see Material and Methods). MYM and MMY lines show intensities in the same range as the high expressing M3GM++ and + lines (Fig. 2B,C). The weaker expressing lines M3GM- and M3GM- -, however, showed significantly lower intensities (Fig. 2B,C).

We collected the 3^rd^ leaf of soil-grown plants 19-21 days after germination and quantified the capacity of the six different complementation lines (MYM, MMY and four M3GM) to rescue the GC abortion and SC recruitment phenotypes of *sid/bdmute-1* (Fig. 2E, Fig. S2). We used four different phenotypic classes: if the GCs showed an oblique division and/or are aborted in their development they were classified as “aborted”. Longitudinally divided, non-aborted GCs that formed a pore were classified according to the number of SCs they recruited (i.e. 0 SC, 1 SC or 2 SC; Fig. 1C, Fig. 2E, Fig. S2). As shown previously, MYM fully complemented both the GC and SC phenotypes, while the mobile, yet non-functional MMY construct failed to complement SC recruitment (Fig. 2E, Fig. S2). In the 3^rd^ leaf, though, MMY seemed to partially complement the GC abortion phenotype (Fig. 2E, Fig. S2), but this was not confirmed in adult leaves (Fig. S8A,B). We, therefore, continued to assume MMY to be non-functional both in terms of SC recruitment and GMC division.

Unexpectedly, we found a dosage-dependent, gradual complementation of both the GC and SC phenotype using the putatively immobile 3xGFP complementation lines (Fig. 2E, Fig. S2). The weakest line (M3GM- -) complemented the GC abortion phenotype almost completely (25.2% ± 4.4% aborted GCs in *sid/bdmute-1* to 2.5% ± 2.2% aborted GCs in M3GM- -) but failed to induce additional SCs (Fig. 2E, Fig. S2). The lines with higher expression (M3GM-, M3GM+) showed a gradual increase in their potential to rescue SC recruitment with over half of the complexes recruiting at least one SC in the M3GM-line and 68.9% ±38.8% of the complexes recruiting two SCs in the M3GM+ lines (Fig. 2E, Fig. S2). Strikingly, the strongest line (M3GM++) almost fully rescued the *sid/bdmute-1* phenotype with the mature leaf displaying 98.5% ± 0.8% WT stomata with two SCs, which was statistically indistinguishable from the MYM full complementation lines, where 99.4% ± 0.7% of stomata recruited 2 SCs (Fig. 2E, Fig. S2). Importantly, this dosage-dependent rescue was confirmed in adult leaves used for gas exchange analysis (see below and Fig. S8).

In conclusion, 3xGFP-BdMUTE was able to rescue the GC and SC phenotypes of *sid/bdmute-1* in a dose-dependent manner.

### Reduced mobility of 3xGFP-BdMUTE leads to delayed subsidiary cell recruitment

The dosage-dependent rescue of SC recruitment by mobility-impaired 3xGFP-BdMUTE implied that BdMUTE mobility might not be required to induce SC identity and divisions. However, careful, quantitative confocal imaging of the strongest 3xGFP-BdMUTE line (M3GM++) revealed a weak 3xGFP-BdMUTE signal in SMCs (Fig. 3A). This suggested that MUTE mobility is not completely abolished but merely reduced by a 3xGFP tag observed. To quantify BdMUTE mobility, we imaged the two lines MYM and M3GM++, which both seemed to fully complement the GC and SC phenotype. We used a 515 nm laser with 3.56% intensity and 75% signal gain for the YFP line and a 489 nm laser with 37.4% intensity and 200% signal gain for the 3xGFP line to visualize signal in the SMC nuclei without achieving signal saturation in the GMC cytoplasm (Fig. 3A). However, GMC nuclear signal was saturated under these conditions (Fig. 3A). We then measured the integrated density of signal in a constant area in the GMC cytoplasm and the SMC nuclei and used the ratio of these intensities as a proxy for BdMUTE mobility from the GC to the SC lineage (Fig. 3B). The GMC cytoplasm to SMC nuclei signal ratio was twice as high for YFP-BdMUTE (0.403 ± 0.151) compared to 3xGFP-BdMUTE (0.216 ± 0.097; Fig. 3B). This suggested that BdMUTE mobility–albeit significantly reduced–was not completely abolished when a 3xGFP tag was added (Fig. 3B).

Even though the mature leaf epidermis of M3GM++ lines suggested a full rescue, we observed a significant delay of SC recruitment in these lines (Fig. 3C,D). In complementing MYM lines, 100% of dividing GMCs have successfully recruited both SCs (Fig. 3C,D). In M3GM++ lines, however, many dividing GMCs recruited only one or no SCs yet (Fig. 3C,D). This suggested that the strict spatiotemporal order of GC and SC divisions, where the SC divisions always precede the symmetric GMC divisions, are severely disturbed when BdMUTE mobility is impaired. To quantify the observed delay in M3GM++ compared to MYM we used two different approaches. First, we analyzed up to 15 developing stomatal complexes apically (i.e. older) and basally (i.e. younger) relative to the first GMC division in different stomatal files of MYM and M3GM++ (Fig. 3D) and scored if these GMCs had already recruited one or two SCs. In all analyzed stomatal files in MYM, 100% of the cells above and below the first GMC division were already flanked by two SCs (Fig. 3D). In M3GM++ on the other hand, only 43% of the cells (n = 89/205 GMCs) had already recruited two SCs, while 26% (n = 54/205 GMCs) had only recruited one SC and 30% (n = 62/205 GMCs) did not recruit any SCs (Fig. 3D). The fraction of GMCs that completed the symmetric division apically to the first GMC division, however, was similar between the two lines (n = 73/94 GMCs = 78% in MYM; n = 81/118 GMCs = 69% in M3GM++; Fig. 3D). Second, the length/width ratio of GMCs is a good proxy for the specific developmental stage between stage 3 and stage 5 (Fig. 1F) (Zhang et al., 2022). We, therefore, measured the length/ width ratio of GMCs and correlated it to the SC recruitment status (0, 1 or 2 SC; Fig. 3E) and GMC division (yes or no; Fig. S3). GMC length/width ratio before GMC division was not different between MYM and M3GM++ (Fig. S3) suggesting that GC development was not disturbed. For SC recruitment, however, only MYM lines showed SC recruitment to be tightly regulated in time and space (Fig. S3E). GMCs in MYM that only recruited one SC showed a very narrow window of GMC length/width ratio (1.06 ± 0.09) and were not very frequent (9%; Fig. 3E). GMCs with a lower length/ width ratio (< 1.0) recruited no SCs yet, and GMCs with a higher length/width ratio (> 1.2) recruited two SCs already (Fig. 3E). In M3GM++, the GMCs that recruited none or only one SC show a wide distribution of length/width ratios (0.98 ± 0.42 and 1.38 ± 0.52 respectively in M3GM++ compared to 0.76 ± 0.20 and 1.06 ± 0.09 in MYM, Fig. 3E). This indicated that the correlation between GMC length/width ratio and number of recruited SCs was severely disturbed in M3GM++. This suggests a significant SC recruitment delay when BdMUTE mobility is impaired. However, even reduced BdMUTE mobility seemed sufficient to eventually induce SMC identity and trigger SC division. This might also explain the strict dosage-dependency of 3xGFP-BdMUTE in its ability to complement the SC recruitment defects; weakly expressed lines might simply not be able to reach sufficient BdMUTE levels in due time in neighboring cells to establish SMC identity.

Together, the spatiotemporal disconnect between GC and SC development when BdMUTE mobility is reduced reinforces the concept of BdMUTE mobility being required to establish SMCs and induce SC divisions. While we could not completely abolish BdMUTE mobility using a bulky 3xGFP tag, the reduction of BdMUTE mobility was sufficient to severely disturb and delay SC recruitment.

### A cell-autonomous role for BdMUTE in guard cell division orientation independent of subsidiary cell recruitment

Another implication of the dosage-dependent 3xGFP-BdMUTE complementation is that weak 3xGFP-BdMUTE expression in the GMC is sufficient to rescue GC abortion irrespective of successful SC recruitment. While GC abortion is almost completely rescued in M3GM- - (Fig. 2E, Fig. S2), this line never formed two SCs and only weakly increased the frequency of single SC complexes from 4.4% ± 0.5% in *sid/bdmute-1* to 8.4% ± 2.7% in M3GM- -. Consequently, the relevant aspect in rescuing the GMC division phenotype seemed to be BdMUTE presence in the GMC (Fig. 2D,E, Fig. S2). This suggested that proper GMC division and formation of functional GC complexes did not depend on successful SC recruitment, that GC and SC divisions are two independent processes, and that *BdMUTE* has a cell-autonomous role during GC formation.

Many of the aborted complexes in *sid/bdmute-1* showed skewed and oblique longitudinal cell divisions, which could also be completely transverse (Fig. 4A). Therefore, *BdMUTE* might have a function in orienting GMC divisions and that wrongly specified division planes and a non-symmetric longitudinal division might cause GC abortion. We thus quantified GMC cell division orientations in WT, *sid/bdmute-1* and *sid/ bdmute-1* complemented with the mobile, yet non-functional MMY and weakest non-mobile M3GM- - line (Fig. 4, Fig. S4). We imaged developmental leaf zones of these four lines right after the GMC division (early stage 5, Fig. 1D) and measured the distances of the newly formed central cell wall to lateral GC walls (Fig. 4B) both on the apical and basal side of the complex (Fig. S4). We then calculated the distance ratio of the shorter and longer distance in all four lines (Fig. 4B, Fig. S4). A ratio of 1 would indicate a perfectly symmetrical division, a ratio smaller than 1 a skewed division. Completely transverse divisions were classified as ratio equals 0 (Fig. S4). We assumed that one side exhibiting skewed division may be sufficient to cause GC complex abortion, therefore we added the top and bottom ratios to investigate the “total skewness” per complex (Fig. 4C,D). Accordingly, a ratio of 2 would represent perfectly symmetrical divisions. Stage 5 GC complexes in WT all show a “total skewness” ratio range of 1.6 - 2 (Fig. 4C). As no stage 5 GC complexes in WT abort, we defined this range as sufficiently symmetric to form functional GC complexes (Fig. 4C, gray dots). Many stage 5 GC complexes showed lower “total skewness” ratios in *sid/bdmute-1* (n = 26/82 complexes below 1.6; 31.7%) and *sid/bdmute-1* transformed with MMY (n = 37/78; 47.4%). In M3GM- -, however, the “total skewness” ratio was almost fully rescued with only 9.1% (6 of 66) of stage 5 GC complexes showing a ratio smaller than 1.6 and none smaller than 1.5 (Fig. 4C).

In conclusion, our data suggested that weak expression of 3xGFP-BdMUTE in GC is sufficient to cell-autonomously rescue division plane orientation and that this process does not require SC recruitment. Furthermore, division plane orientation defects seem indeed to be strongly correlated with GC complex abortion in *sid/bdmute-1* and thus potentially causative for failed GC development in grass *mute* mutants. However, more GMCs display a skewness ratio smaller than 1.6 than actually abort. This suggests that slightly skewed GMC divisions can nonetheless result in functional GC complexes.

### Dumbbell guard cell morphogenesis requires subsidiary cell recruitment

So far, our data suggested that the division programs of the GC and SC lineage are largely independent of one another with BdMUTE controlling two independent aspects in the respective lineages. On the one hand, a reduction in BdMUTE mobility through the addition of a bulky 3xGFP tag delayed SC recruitment significantly suggesting that mobility was required to non-cell-autonomously induce SC divisions within a flexible, broad window of opportunity. On the other hand, BdMUTE is cell-autonomously required in GMCs for division symmetry and division orientation and these two processes are mostly independent of whether SCs are recruited or not. Yet, GCs that are or are not flanked by SCs displayed distinct morphologies and are stockier and rounder with a less pronounced dumbbell shape (Fig. 5). GCs flanked by SCs, on the other hand, displayed the characteristic dumbbell shape of grass GCs (Fig. 5). To quantify this, we used mature leaves of the M3GM-line that showed the largest diversity of SC recruitment phenotypes with almost equal parts recruiting zero, one or two SCs (Fig. 2E, Fig. S2, Fig. S8). We quantified total GC length and GC width at apices and in the central rods of stomatal complexes with zero, one and two SCs (Fig. 5A,B; Fig. S5). Stomatal complexes with two SCs were significantly longer than those without or with only one SC, which showed intermediate GC lengths (Fig. S5). To describe dumbbell morphology, we calculated the apex-to-middle ratio as a proxy for curvature of the GCs and found a significant difference between GCs with and without SCs (Fig. 5A,B). In a stomatal complex without SCs, the GCs had a lower mean apex/middle ratio (1.10 ± 0.13) than GCs in a stomatal complex with two SCs (1.56 ± 0.24; Fig. 5B). This suggested that morphogenesis either depended on the presence of a flanking SC or different levels of functional BdMUTE protein in GMCs and a cell-autonomous role of BdMUTE in GC morphogenesis. Yet, even within a single complex that only recruited one SC the morphology of the two GCs was starkly different; the GC flanked by an SC showed a higher apex/middle ratio (1.30 ± 0.19) compared to the non-flanked GC (1.11 ± 0.20; Fig. 5B). In these cases, both GCs stem from the same GMC, and hence likely inherited the same amount of BdMUTE protein. The distinguishing factor was only the presence or absence of a flanking SC. This strongly suggested a non-cell-autonomous effect of the SCs on GC morphogenesis. However, a single SC was not sufficient to fully reconstitute the apex/middle ratio of the flanked GC (Fig. 5B) nor the GC length (Fig. S5). The GC length of complexes with a single SC (21.4 μm ± 1.64 μm) was between complexes with two SCs (24.7 μm ± 2.59 μm) or no SCs (19.6 μm ± 2.01 μm; Fig. S5). This indicates that GC elongation and GC morphogenesis both depend on successful recruitment of two lateral SCs and that they might be functionally linked.

An issue of using brightfield DIC images to quantify apex/middle ratios of GCs is that the focal plane of individual stoma is difficult to control, potentially introducing a measurement bias. We therefore generated 3D confocal stacks of individual stomata with no, one or two SCs in M3GM- (Fig. 5E-G). We then quantified the GC area of optical XZ cross-sections of the central rod and the two apices of M3GM-complexes with zero, one and two SCs (Fig. 5C,D). GCs that are not flanked by SCs show a consistent mean GC circumference at the apex/middle ratio irrespective of whether they are part of a stomatal complex that recruited no or one SC (1.81 ± 0.45 in complexes with 1 SC and 1.86 ± 0.15 in complexes with 0 SC, Fig. 5D). In contrast, GCs that were flanked by a SC in complexes with only one SC showed a higher apex/middle ratio (2.33 ± 0.77; Fig. 5D). However, GCs from wild type-like (in M3GM-) stomatal complexes with two SCs showed the highest apex/middle area ratio (2.86 ± 0.72) and thus the most pronounced dumbbell shape (Fig. 5D).

Together, this suggests that dumbbell morphogenesis is indeed non-cell-autonomously regulated by a flanking SC. Yet, when only one SC is recruited, morphogenesis of the flanked GC is incomplete, which might be related to impaired GC elongation in single SC complexes.

### Gradual complementation of the *sid/bdmute-1* mutant phenotype is quantitatively reflected in leaf-level gas exchange physiology

Previous work showed the importance of SCs to grass stomatal function (Durney et al., 2023; Liu et al., 2024; Raissig et al., 2017; Zhang et al., 2022). In *sid/bdmute-1* plants, absolute stomatal conductance (*g*_sw_), speed of stomatal movements, maximum stomatal aperture and maximum stomatal conductance were drastically impaired (Raissig et al., 2017). To test if gradual rescue of the GC and SC defects in *sid/ bdmute-1* was reflected in stomatal gas exchange parameters, leaf-level gas exchange was investigated under fluctuating light intensities (1000 μmol m^-2^ s^-1^) -> 100 μmol m^-2^ s^-1^ -> 1000 μmol m^-2^ s^-1^ -> 0 μmol m^-2^ s^-1^). For wild type, MYM, MMY, *sid/bdmute-1* and the four M3GM (- - to ++) lines, we measured and calculated *g*_sw_, carbon assimilation (*A*) and intrinsic water-use efficiency (iWUE). As reported previously, *sid/bdmute-1* shows significantly reduced *g*_sw_ and *A* in high light conditions (1000 μmol m^-2^ s^-1^; Fig. 6A,C, Table 1, Fig. S6). These differences are more extreme for *g*_sw_ than *A* (Fig. 6A,C, Table 1, Fig. S6), leading to higher iWUE in *sid/bdmute-1* compared to WT at high light (Fig. 6E,F, Table 1, Fig. S7). Similar observations can be made in functional and complementing MYM versus non-functional and non-complementing MMY lines with the former being very similar to WT and the latter strongly resembling *sid/bdmute-1* (Fig. 6A,C,E; Table 1: Fig. S6; Fig. S7). Strikingly, the dosage-dependent, gradual complementation of stomatal morphology in the M3GM lines is fully mirrored in their respective *g*_sw_, *A* and iWUE (Fig. 6B,D,F, Table 1, Fig. S6, Fig. S7). This strongly confirms the importance of stomatal morphology and presence of SCs on gas exchange in grasses. This is further confirmed by calculating physiological parameters per stoma (Fig. S9) rather than per leaf area (Fig. 6). Stomatal density, despite being slightly different in some lines, does not explain the major significant differences in *g*_sw_ and *A* (Fig. S6, Fig. S8C). Stomatal morphology of the leaves used for gas exchange measurements, however, quantitatively recapitulated the phenotypes observed in the 3^rd^ leaf (Fig. 2E, Fig. S2, Fig. S8A,B). Therefore, our data strongly suggested that stomatal morphology affected gas exchange levels in the assessed lines in a quantitative manner with four-celled WT-like complexes allowing for higher and a much wider quantitative range of *g*_sw_ and *A*.

**Table 1:**
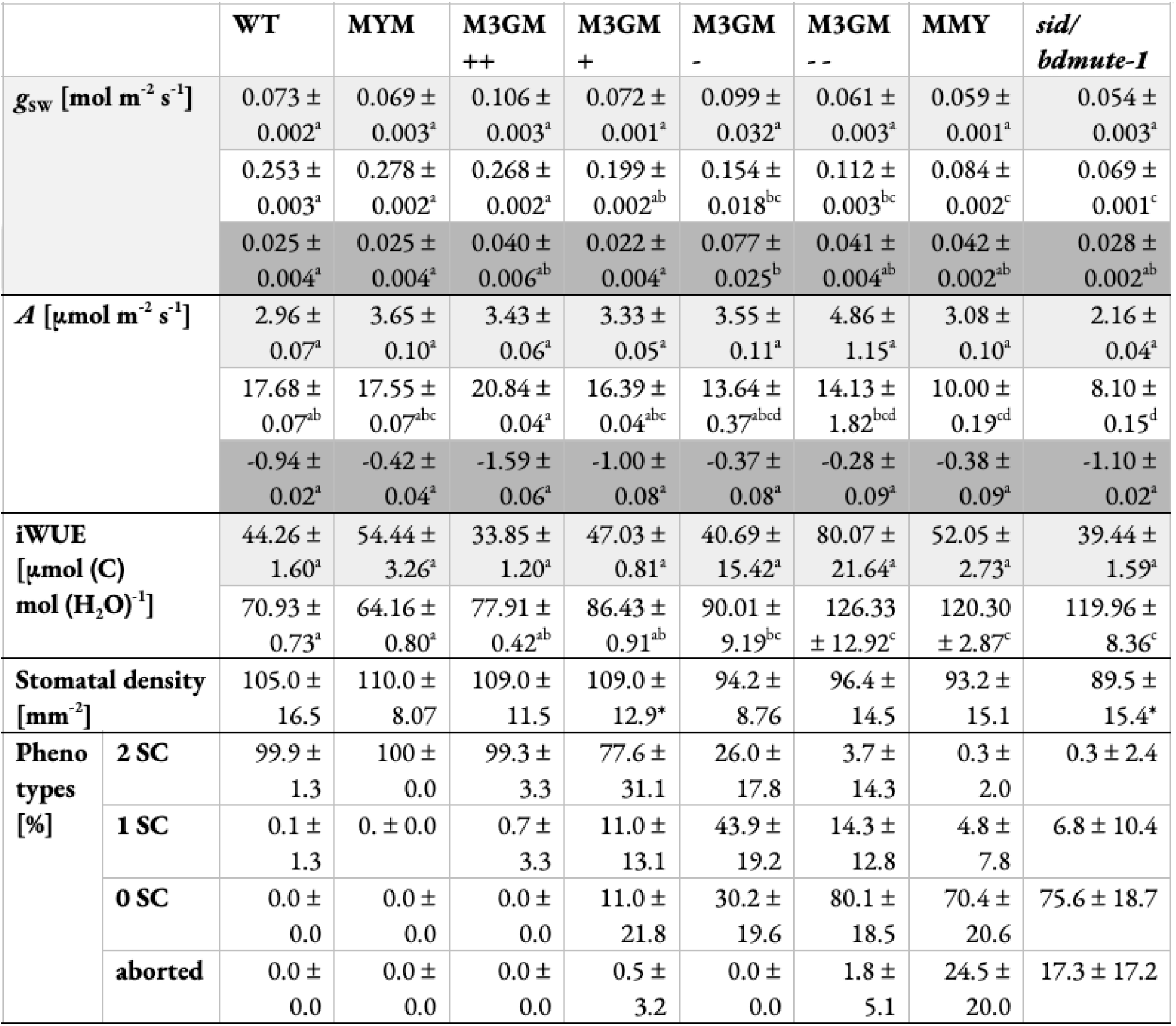
Steady state gas exchange data and stomatal anatomy of analyzed lines. Data relates to Fig. 6 and Fig. S6-9. Grayscale background in g_SW_, A and iWUE indicates light intensity: White = 1000 μmol m^-2^ s^-1^, light gray = 100 μmol m^-2^ s^-1^, dark gray = 0 μmol m^-2^ s^-1^. Significant differences for g_SW_, A and iWUE are indicated with differing superscript letters. Statistical test: ANOVA followed by Tukey’s HSD test (alpha = 0.05). Significant differences to the WT individuals investigated in the same experimental run (n = 4-6) for stomatal density are indicated with asterisks. Statistical test: Two-sided Student’s t-test (significant if p < 0.05).

**Figure 6:**
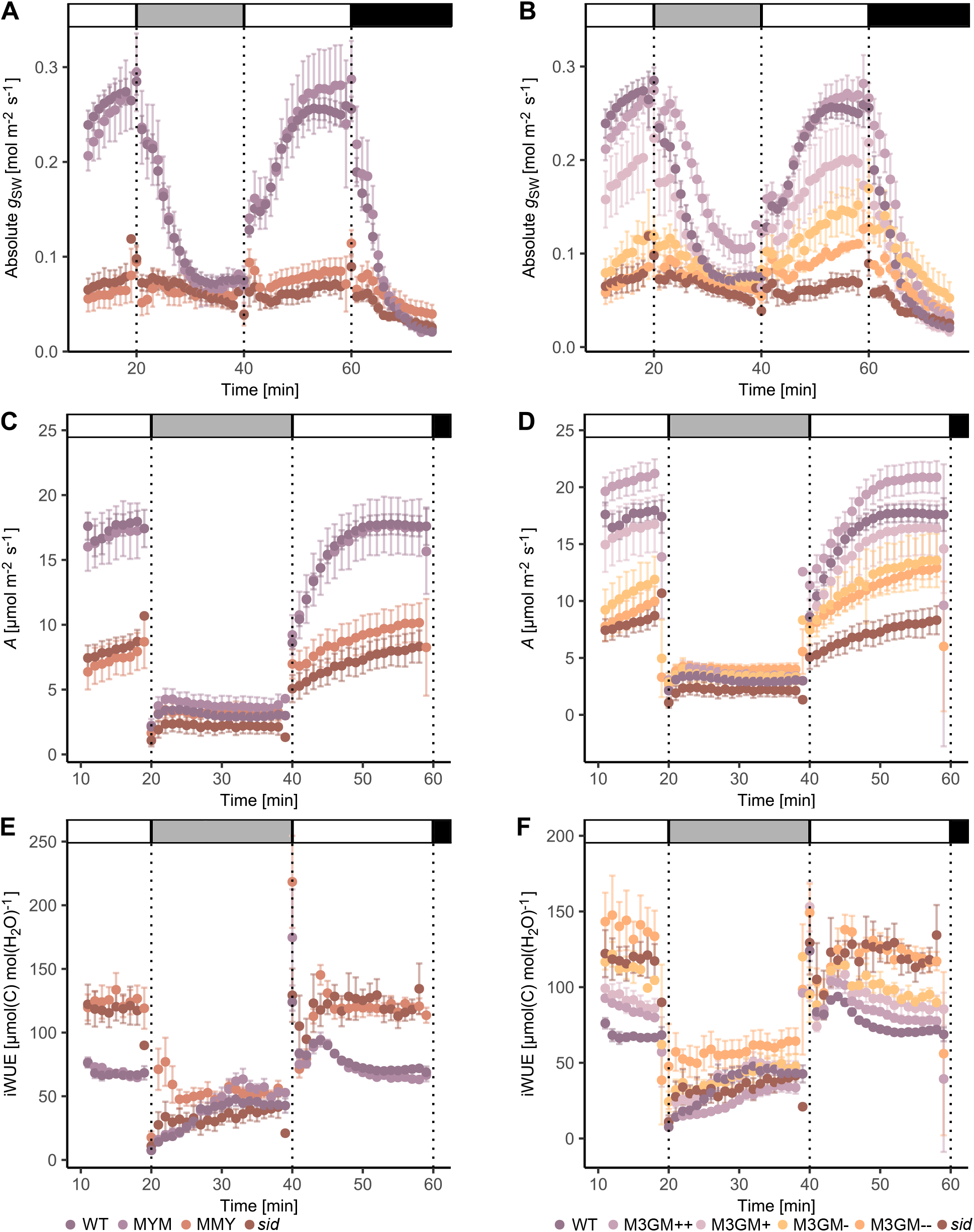
Physiological complementation follows cytological complementation. (**A**) Absolute stomatal conductance (*g*_SW_) of wild type (WT), *sid/bdmute-1*;*BdMUTEp:YFP-BdMUTE* (MYM), *sid/bdmute-1*;*BdMUTEp:BdMUTE-YFP* (MMY) and *sid/bdmute-1* under changing light conditions as indicated by grayscale bars above each plot: White = 1000 μmol m^-2^ s^-1^, light gray = 100 μmol m^-2^ s^-1^, black = 0 μmol m^-2^ s^-1^. (**B**) *g*_SW_ of WT, the four *sid/ bdmute-1*;*BdMUTEp:3xGFP-BdMUTE* (M3GM) lines (++, +, - and - -) and *sid/bdmute-1* under changing light conditions as indicated by grayscale bars. (**C**) Carbon assimilation (*A*) of WT, MYM, MMY and *sid/bdmute-1* under changing light conditions as indicated by grayscale bars. (**D**) *A* of WT, M3GM lines and *sid/bdmute-1* under changing light conditions as indicated by grayscale bars. (**E**) Intrinsic water-use efficiency (iWUE) of WT, MYM, MMY and *sid/bdmute-1* under changing light conditions as indicated by grayscale bars. (**F**) iWUE of WT, M3GM lines and *sid/bdmute-1* under changing light conditions as indicated by grayscale bars. Note that the same WT and *sid/bdmute-1* data is shown in A, C, E and B, D, F. Measured were the youngest, fully expanded leaves of 3-4 week old, soil-grown plants; n = 5-6 individuals per genotype, 26 individuals for WT. Dots represent the means of all individuals and error bars indicate standard error. Data depicted is used to calculate the steady state values in Table 1, Fig. S6 and Fig. S7. The adult leaves assessed for gas exchange were also used to quantify the complementation of *sid/bdmute-1*-induced stomatal morphological defects of the different complementation lines and to calculate stomatal densities (Table 1, Fig. S8).

## DISCUSSION

In this study, we addressed open questions regarding BdMUTE’s function during SC and GC formation in grasses (Fig. 1). We were able to show that reducing BdMUTE mobility resulted in spatiotemporally delayed SC recruitment indicating that BdMUTE mobility is indeed required to recruit SCs (Fig. 3). In addition, BdMUTE guides GC division orientation in a cell-autonomous manner (Fig. 4) that does not require SC recruitment (Fig. 2). However, complete morphogenesis of dumbbell GCs requires bilateral SC recruitment (Fig. 5). Finally, the dose-dependent rescue of GC and SC formation in *sid/bdmute-1* by 3xGFP-BdMUTE (Fig. 2) was mirrored in a gradual complementation of stomatal physiology (Fig. 6). The distinct origin of GCs and SCs and the strict developmental gradient of stomatal development in the model grass *Brachypodium distachyon* (purple false brome; Pooideae) allowed us to further untangle the complex, multifaceted roles of BdMUTE during grass stomatal formation.

### Subsidiary cell recruitment is linked to MUTE mobility

MUTE protein was shown to be a mobile factor (Raissig et al., 2017; Wang et al., 2019). In maize, rice and *B. distachyon*, MUTE is expressed in GMCs and subsequently moves laterally into the two neighboring cell files (Raissig et al., 2017; Wang et al., 2019). While lateral deployment of a core stomatal transcription factor through cell-to-cell mobility to induce “ectopic” stomatal fate is a compelling hypothesis, it still lacked experimental proof. We attempted to make an immobile version by tagging BdMUTE with 3xGFP that has been shown to hinder protein mobility (Andersen et al., 2018; Clark et al., 2016; Goretti et al., 2020) but saw full rescue of *sid/bdmute-1* in lines with high protein dosage. This initially suggested that SC recruitment is not dependent on BdMUTE presence in the lateral cell files, but rather relying on secondary, putatively mobile factors downstream of MUTE like peptides, small RNAs, or plant hormones.

However, careful imaging revealed reduced but not abolished mobility that, within a window of opportunity, was sufficient to reach the required threshold to induce SC divisions (Fig. 3C-E). Other studies reported a steric 3xGFP tag as sufficient to immobilize proteins (Andersen et al., 2018; Clark et al., 2016; Goretti et al., 2020; Lucas et al., 1995). This could be due to a more narrow window of opportunity in which reduced mobility is sufficient to prevent the protein from reaching the required threshold and activate a specific developmental process. Alternatively, plasmodesmata in different tissue contexts or even species might have distinct size exclusion limits affecting the degree of protein mobility impairment by adding 3xGFP (Bayer and Benitez-Alfonso, 2024). Also, different cell types and tissue might express distinct proteins that promote cell-to-cell mobility. The essential protein SHORTROOT INTERACTING EMBRYONIC LETHAL (SIEL), for example, interacts with different mobile proteins like the root patterning transcription factor SHORTROOT (SHR) or the hair cell patterning transcription factor CAPRICE and promotes protein mobility of SHR during root patterning (Koizumi et al., 2011). Finally, we cannot fully exclude cleavage of the 3xGFP fusion protein, which could only be tested with a BdMUTE-specific antibody.

Taken together, our results strongly suggest that the mobile transcription factor BdMUTE must be present in both GMCs and SMCs to spatiotemporally coordinate SC recruitment and GMC division, which are two largely independent processes. Strikingly, we find large temporal flexibility regarding the time point of SC formation. Even severely delayed SC formation in M3GM++ leads to WT stomata in mature leaf zones. However, since GC morphogenesis and complex maturation cannot be fully uncoupled from the presence of SCs, we cannot exclude that delayed SC recruitment can quantitatively affect GC morphogenesis. Even slight disturbances in the GC maturation process could have functional consequences when grown in competition or in harsher and fluctuating environmental conditions.

### BdMUTE guides guard cell division plane orientation in a cell-autonomous manner

In *Arabidopsis*, stomatal patterning relies on the self-renewal of meristemoids that can undergo amplifying divisions before acquiring GMC identity (Bergmann and Sack, 2007; Cheng and Raissig, 2023; Lau and Bergmann, 2012). AtMUTE controls the transition from meristemoid to GMC and directly controls a regulatory network of cell-cycle genes and their repressors, which enables a single symmetric division of the GMC (MacAlister et al. 2007; MacAlister and Bergmann 2011; Han et al. 2018; Pillitteri et al. 2007; Han et al. 2022; Kim et al. 2022; Weimer et al. 2018). In grasses, stomatal development is restricted to the most basal part of the developing leaf and takes place in specialized cell files (Stebbins and Shah, 1960). The first asymmetric cell division (ACD) directly gives rise to a GMC omitting the meristemoid stage (Raissig et al., 2016).

Thus, division capacity in the grass leaf epidermal lineages is fundamentally different and spatially restricted to the leaf base and to stage 1 and stage 2 stomatal cells (Fig. 1D). SMC and GMC divisions both occur beyond the “transverse division competency zone”. Therefore, GC fate commitment in grasses might not require division competency inhibition like in eudicots but a division competency establishment instead (McKown and Bergmann, 2020; Nunes et al., 2020). Indeed, FAMA proteins in grasses lack the conserved LxCxE motif that mediates the interaction with the cell cycle inhibitor RETINOBLASTOMA RELATED1 (RBR1) involved in repressing division competency in GCs (Matos et al., 2014; Spiegelhalder and Raissig, 2021). Accordingly, grass *fama* stomata arrest after the symmetric GMC division and fail to differentiate a pore and functional GCs but do not show any additional divisions (Liu et al., 2009; McKown et al., 2023; Wu et al., 2019). We can only speculate what establishes and regulates division competency in the grass GC lineage. Clearly, *mute* mutant GMCs can divide in both wild and domesticated grasses, but fail to define division plane orientation (Raissig et al., 2017; Wang et al., 2019; Wu et al., 2019). Potentially, the duplicated SPCH genes regulate division competency both in stage 1 stomatal cells and in GMCs. BdSPCH1, in particular, is strongest expressed in stage 2 and stage 3 GMCs, while BdSPCH2 is strongest expressed in stage 1 stomatal cells (Raissig et al., 2016). Indeed, we observe no impaired cell division capacity in the GC lineage in *sid/bdmute-1*. Even though (functional) stomatal index and stomatal density is reduced in *sid/bdmute-1* (Fig. S10A,C), this reduction completely disappears when also aborted complexes are counted (i.e. stomatal lineage index and density; Fig. S10B,D). This suggests that no division defects in stage 1/stage 2 stomatal cells occur in *sid/bdmute-1*. In the SC lineage, however, the establishment of SMC identity is inherently linked to division competency. Therefore, MUTE does regulate division competency in SMCs as very few SC divisions occur in *mute* mutants (Raissig et al., 2017; Wang et al., 2019; Wu et al., 2019).

Both, the highly asymmetric divisions we see in SMCs and the symmetric divisions of the GMC are unique in the grass epidermis as they are some of the rare longitudinal divisions. Most epidermal divisions in the division zone are transverse; longitudinal divisions are actively inhibited for example by *BROAD LEAF1* in barley (*HvBL1*; Jöst et al., 2016) in young leaf primordia. Grass MUTE protein–being present in both GMCs and SMCs–might be linked to the specification of longitudinal division orientation. Indeed, MUTE expression in GMCs, but not SC recruitment, is required for GMC division orientation (Fig. 2E, Fig. 4, Fig. S2, Fig. S8A,B) as has been previously hypothesized, but not experimentally shown (Serna, 2021). In SMCs, MUTE induces BdPOLAR (Zhang et al., 2022), a distal polarity program that together with a MUTE independent proximal program (Cartwright et al., 2009; Facette et al., 2015; Humphries et al., 2011; Zhang et al., 2012; Zhang et al., 2022) guides the asymmetric cell division (ACD) of SMCs. In particular, the distal BdPOLAR domain, which is absent in *bdmute*, seems to regulate cortical division site orientation linking BdMUTE to a division orientation program in SMCs (Zhang et al., 2022). Yet, BdPOLAR is normally expressed in *bdmute* GMCs suggesting that a distinct program orients division planes in GMCs (Zhang et al., 2022). Specific cyclins, for example, were shown to be involved in both, symmetric cell division (SCD) (Weimer et al., 2018; Han et al., 2018) and ACD (Cruz-Ramírez et al., 2012) during stomatal development and MUTE has been shown to regulate expression of such cell-cycle factors. For example, MUTE upregulates the expression of *CYCA3;2, CYCD5;1* and *CYCD7;1* involved in the *Arabidopsis* GC SCD (Han et al., 2018; Weimer et al., 2018; Zuch et al., 2023). In cortex-endodermal initial cells in the root ground tissue, CYCD6;1 is specifically activated and required for normal formative and patterning division (Sozzani et al., 2010). Strikingly, proper formation of cortex and endodermis also involves a division plane reorientation, where upon a formative transverse division a periclinal division gives rise to an endodermal and a cortex cell (Dolan et al., 1993). Therefore, we propose a grass GMC-specific cell division program, in which the SPCHs establish division capacity and MUTE controls division orientation.

It remains unclear, though, why in domesticated grasses, *mute* mutants completely fail to execute proper GC divisions (Wang et al., 2019; Wu et al., 2019). In *zmmute*, for example, GCs can undergo several rounds of transversal divisions (Wang et al., 2019). Possibly, a domestication-induced loss of genetic diversity and, concomitantly, a loss of a developmental buffering system could underlie this severe and seedling-lethal phenotype in maize and rice. Alternatively, the developmental leaf zone in *B. distachyon* is much smaller than in maize and even rice. Therefore, *MUTE* and *FAMA* expression windows overlap significantly. In fact, BdFAMA has been shown to partially rescue the aborted division phenotype in *sid/bdmute-1* (McKown et al., 2023), particularly when it is expressed under the *BdMUTE* promoter. In a mutant background, where either MUTE or FAMA is broken, the functional homolog can bind the same heterodimerization partners and could sufficiently activate the relevant target genes. Without the *MUTE* and *FAMA* expression windows overlapping due to longer developmental zones in rice and particularly in maize, however, FAMA might not be able to sufficiently substitute for MUTE.

### Subsidiary cells influence guard cell morphogenesis

In *sid/bdmute-1*, GCs are shorter and stockier with less pronounced dumbbell shape (Fig. 1C, Fig. 5, Fig. S5). We observe partial reconstitution of the GC shape (apex/middle ratio) and length in stomatal complexes with just one SC, yet only those GCs that were actually flanked by the SC reconstituted shape (Fig. 5). However, GC morphogenesis and elongation is only complete, when both GCs are flanked by SCs. This suggests that the presence of lateral SCs non-cell autonomously influences GC shape independent of MUTE protein levels in GMCs. Whether this involves cell-to-cell signaling processes, biomechanical processes and/or the specific maturation processes of the shared apoplast between GCs and SCs remains yet to be determined. In mature grass stomata, GCs and SCs are thought to oppositely shuffle osmolytes and thus execute opposite turgor pressures facilitating rapid opening and closing kinetics (Durney et al., 2023; Franks and Farquhar, 2007). Potentially, active pressurization of the SCs during stomatal development already might support GC elongation and morphogenesis. Biomechanical processes were repeatedly shown to influence plant cell morphogenesis in general and in GCs in particular (Elliott et al., 2024; Geitmann and Ortega, 2009; Landrein and Hamant, 2013; Rui et al., 2019; Yi et al., 2019). Therefore, mechanical influence of the SCs seems likely to be at least partially involved in guiding the morphogenesis of GCs in grasses.

In GCs without a SC we see a thicker middle rod with more pronounced cell wall staining, which indicates excessive cell wall formation (Fig 5E). This could be a byproduct of less elongated GCs and consequently, a non-elongated surplus of cell wall material. In stomatal complexes with a single SC, however, both GCs have the same length, but we still primarily observe a stronger cell wall signal in GCs that were not flanked by SCs. Thus, cell wall build up might indicate a form of compensatory effect, possibly induced by altered biomechanical interactions with flanking pavement cells instead of flanking SCs.

### The dose-dependent cytological rescue is mirrored by a dose-dependent physiological rescue

Different levels of 3xGFP-MUTE caused a gradual, dosage-dependent rescue of both GC and SC division phenotypes. Physiological measurements showed that this cytological complementation gradient is fully reflected in gradual rescue of the stomatal physiological defects of *sid/bdmute-1* (Fig. 6, Fig. S6-9).

Stomatal density was shown to affect leaf-level gas exchange in wild type *B. distachyon* (Nunes et al. 2022) and transgenic cereal crops (Caine et al., 2018; Dunn et al., 2019; Hughes et al., 2017; Karavolias et al., 2023). Out of all lines included in this study, only *sid/bdmute-1* showed a significantly decreased stomatal density in comparison to the WT plants grown at the same time (Fig. S8C) suggesting that the gradually lower *g*_SW_ and *A* under high light conditions in the M3GM lines was unlikely due to this anatomical trait (Fig. 6A-D, Fig. S6). In addition, the gradual differences remained even when the gas exchange parameters were calculated per stoma (Fig. S9), which further indicated a much more dominant role of stomatal anatomy rather than density on gas exchange.

A recent study on maize *pangloss* mutants hypothesized that SCs are required for full stomatal closure as stomatal complexes with defective SCs have increased pore aperture in comparison to WT even under closure-inducing conditions (Liu et al., 2024). The strongest differences we observed were for stomatal aperture (indicated by *g*_SW_) of complexes lacking SCs under high light conditions (1000 μmol m^-2^ s^-1^) (Fig. 6, Fig. S6A,C,E). Low (100 μmol m^-2^ s^-1^) or no light (0 μmol m^-2^ s^-1^) treatments showed mostly non-significant albeit observable effects on *g*_SW_ and *A* in the M3GM lines, MMY, and *sid/ bdmute-1* (Fig. 6, Fig. S6). This suggests that presence or absence of SCs predominantly affects stomatal opening but also stomatal closing in *B. distachyon*.

Moreover, we observed that the presence or absence of SCs in the M3GM lines influenced GC shape and length (Fig. 3E, Fig. 5, Fig. S5). As stomatal size was already shown to affect stomatal physiology in *B. distachyon* (Nunes et al., 2022), we are currently unable to completely untangle SC presence from GC morphology and size. Therefore, it remains unclear if the decreased stomatal performance in the lines described here is due to the lack of SCs and defective ion shuffling and reciprocal turgor regulation or is caused by altered GC morphology and size. To dissect GC form from SC presence, we would require a mutant, which affects SC function rather than SC formation (i.e. *bdmute*, or *bdpolar;bdpan1*).

Nonetheless, SC presence and GC morphology and size likely regulate the rapid stomatal movements in the grass family in a highly synergistic and complementary manner (Franks and Farquhar 2007, McAusland et al., 2016, Nunes et al., 2020).

## ACKNOWLEDGEMENT

We would like to thank the research gardeners Michael Schilbach (COS Heidelberg), Jasmin Sekulovski, Sarah Dolder and Christopher Ball (all three University of Bern). We would like to thank Jan Lohmann and Thomas Greb for discussions and insights, Jan Lohmann for providing reagents (3xGFP cloning vector), and Annika Guse for access to the SP8 confocal microscope.

## REFERENCES

Andersen, T. G., Naseer, S., Ursache, R., Wybouw, B., Smet, W., De Rybel, B., Vermeer, J. E. M. and Geldner, N. (2018). Diffusible repression of cytokinin signalling produces endodermal symmetry and passage cells. Nature 555, 529–533.

Bayer, E. M. and Benitez-Alfonso, Y. (2024). Plasmodesmata: Channels Under Pressure. Annu. Rev. Plant Biol.

Berg, L. S. (2024). licornetics: Plot and calculate physiological parameters using LI-COR data.

Bergmann, D. C. and Sack, F. D. (2007). Stomatal development. Annu. Rev. Plant Biol. 58, 163–181.

Berry, J. A., Beerling, D. J. and Franks, P. J. (2010). Stomata: key players in the earth system, past and present. Curr. Opin. Plant Biol. 13, 233–240.

Caine, R. S., Yin, X., Sloan, J., Harrison, E. L., Mohammed, U., Fulton, T., Biswal, A. K., Dionora, J., Chater, C. C., Coe, R. A., et al. (2018). Rice with reduced stomatal density conserves water and has improved drought tolerance under future climate conditions. New Phytol.

Cartwright, H. N., Humphries, J. A. and Smith, L. G. (2009). PAN1: a receptor-like protein that promotes polarization of an asymmetric cell division in maize. Science 323, 649–651.

Chang, G., Ma, J., Wang, S., Tang, M., Zhang, B., Ma, Y., Li, L., Sun, G., Dong, S., Liu, Y., et al. (2023). Liverwort bHLH transcription factors and the origin of stomata in plants. Curr. Biol.

Chater, C., Kamisugi, Y., Movahedi, M., Fleming, A., Cuming, A. C., Gray, J. E. and Beerling, D. J. (2011). Regulatory mechanism controlling stomatal behavior conserved across 400 million years of land plant evolution. Curr. Biol. 21, 1025–1029.

Cheng, X. and Raissig, M. T. (2023). From grasses to succulents - development and function of distinct stomatal subsidiary cells. New Phytol. 239, 47–53.

Clark, N. M., Hinde, E., Winter, C. M., Fisher, A. P., Crosti, G., Blilou, I., Gratton, E., Benfey, P. N. and Sozzani, R. (2016). Tracking transcription factor mobility and interaction in Arabidopsis roots with fluorescence correlation spectroscopy. Elife 5, 78.

Clark, J. W., Harris, B. J., Hetherington, A. J., Hurtado-Castano, N., Brench, R. A., Casson, S., Williams, T. A., Gray, J. E. and Hetherington, A. M. (2022). The origin and evolution of stomata. Curr. Biol. 32, R539–R553.

Cruz-Ramírez, A., Díaz-Triviño, S., Blilou, I., Grieneisen, V. A., Sozzani, R., Zamioudis, C., Miskolczi, P., Nieuwland, J., Benjamins, R., Dhonukshe, P., et al. (2012). A Bistable Circuit Involving SCARECROW-RETINOBLASTOMA Integrates Cues to Inform Asymmetric Stem Cell Division. Cell.

de Mendiburu, F. (2023). agricolae: Statistical Procedures for Agricultural Research.

Dolan, L., Janmaat, K., Willemsen, V., Linstead, P., Poethig, S., Roberts, K. and Scheres, B. (1993). Cellular organisation of the Arabidopsis thaliana root. Development 119, 71–84.

Dunn, J., Hunt, L., Afsharinafar, M., Al Meselmani, M., Mitchell, A., Howells, R., Wallington, E., Fleming, A. J. and Gray, J. E. (2019). Reduced stomatal density in bread wheat leads to increased water-use efficiency. J. Exp. Bot.

Durney, C. H., Wilson, M. J., McGregor, S., Armand, J., Smith, R. S., Gray, J. E., Morris, R. J. and Fleming, A. J. (2023). Grasses exploit geometry to achieve improved guard cell dynamics. Curr. Biol.

Edwards, D., Kerp, H. and Hass, H. (1998). Stomata in early land plants: an anatomical and ecophysiological approach. J. Exp. Bot. 49, 255–278.

Elliott, L., Kalde, M., Schürholz, A.-K., Zhang, X., Wolf, S., Moore, I. and Kirchhelle, C. (2024). A self-regulatory cell-wall-sensing module at cell edges controls plant growth. Nat Plants.

El-Sharkawey, A. E. (2016). Calculate the Corrected Total Cell Fluorescence (CTCF).

Facette, M. R., Park, Y., Sutimantanapi, D., Luo, A., Cartwright, H. N., Yang, B., Bennett, E. J., Sylvester, A. W. and Smith, L. G. (2015). The SCAR/WAVE complex polarizes PAN receptors and promotes division asymmetry in maize. Nat Plants 1, 14024.

Franks, P. J. and Farquhar, G. D. (2007). The mechanical diversity of stomata and its significance in gas-exchange control. Plant Physiol. 143, 78–87.

Galatis, B. and Apostolakos, P. (2004). The role of the cytoskeleton in the morphogenesis and function of stomatal complexes. New Phytol. 161, 613–639.

Geitmann, A. and Ortega, J. K. E. (2009). Mechanics and modeling of plant cell growth. Trends Plant Sci. 14, 467–478.

Goretti, D., Silvestre, M., Collani, S., Langenecker, T., Méndez, C., Madueño, F. and Schmid, M. (2020). TERMINAL FLOWER1 Functions as a Mobile Transcriptional Cofactor in the Shoot Apical Meristem. Plant Physiol. 182, 2081–2095.

Gray, A., Liu, L. and Facette, M. (2020). Flanking Support: How Subsidiary Cells Contribute to Stomatal Form and Function. Front. Plant Sci. 11, 881.

Haas, A. and Raissig, M. (2020). Seed sterilization and seedling growth on plates in the model grass Brachypodium distachyon. Bio Protoc. 10, e3700–e3700.

Han, S.-K., Qi, X., Sugihara, K., Dang, J. H., Endo, T. A., Miller, K. L., Kim, E.-D., Miura, T. and Torii, K. U. (2018). MUTE Directly Orchestrates Cell-State Switch and the Single Symmetric Division to Create Stomata. Dev. Cell 45, 303–315.e5.

Han, S.-K., Herrmann, A., Yang, J., Iwasaki, R., Sakamoto, T., Desvoyes, B., Kimura, S., Gutierrez, C., Kim, E.-D. and Torii, K. U. (2022). Deceleration of the cell cycle underpins a switch from proliferative to terminal divisions in plant stomatal lineage. Dev. Cell 57, 569–582.e6.

Harris, B. J., Harrison, C. J., Hetherington, A. M. and Williams, T. A. (2020). Phylogenomic Evidence for the Monophyly of Bryophytes and the Reductive Evolution of Stomata. Curr. Biol. 30, 2001–2012.e2.

Hetherington, A. M. and Woodward, F. I. (2003). The role of stomata in sensing and driving environmental change. Nature 424, 901–908.

Hughes, J., Hepworth, C., Dutton, C., Dunn, J. A., Hunt, L., Stephens, J., Waugh, R., Cameron, D. D. and Gray, J. E. (2017). Reducing Stomatal Density in Barley Improves Drought Tolerance without Impacting on Yield. Plant Physiol. 174, 776– 787.

Humphries, J. A., Vejlupkova, Z., Luo, A., Meeley, R. B., Sylvester, A. W., Fowler, J. E. and Smith, L. G. (2011). ROP GTPases act with the receptor-like protein PAN1 to polarize asymmetric cell division in maize. Plant Cell 23, 2273–2284.

Jones, V. A. S. and Dolan, L. (2017). MpWIP regulates air pore complex development in the liverwort Marchantia polymorpha. Development 144, 1472–1476.

Jöst, M., Hensel, G., Kappel, C., Druka, A., Sicard, A., Hohmann, U., Beier, S., Himmelbach, A., Waugh, R., Kumlehn, J., et al. (2016). The INDETERMINATE DOMAIN Protein BROAD LEAF1 Limits Barley Leaf Width by Restricting Lateral Proliferation. Curr. Biol. 26, 903–909.

Karavolias, N. G., Patel-Tupper, D., Seong, K., Tjahjadi, M., Gueorguieva, G.-A., Tanaka, J., Gallegos Cruz, A., Liebermann, S., Litvak, L., Dahlbeck, D., et al. (2023). Paralog editing tunes rice stomatal density to maintain photosynthesis and improve drought tolerance. Plant Physiol.

Kassambara, A. (2023). ggpubr: “ggplot2” Based Publication Ready Plots.

Kellogg, E. A. (2001). Evolutionary history of the grasses. Plant Physiol. 125, 1198–1205.

Kellogg, E. A. (2015). Brachypodium distachyon as a Genetic Model System. Annu. Rev. Genet. 49, 1–20.

Kim, E.-D., Dorrity, M. W., Fitzgerald, B. A., Seo, H., Sepuru, K. M., Queitsch C., Mitsuda, N., Han, S.-K. and Torii, U. (2022). Dynamic chromatin accessibility deploys heterotypic cis/trans-acting factors driving stomatal cell-fate commitment. Nat Plants 8, 1453–1466.

Koizumi, K., Wu, S., MacRae-Crerar, A. and Gallagher, K. (2011). An essential protein that interacts with endosomes and promotes movement of the SHORT-ROOT transcription factor. Curr. Biol. 21, 1559–1564.

Kurihara, D., Mizuta, Y., Sato, Y. and Higashiyama, T. (2015). ClearSee: a rapid optical clearing reagent for whole-plant fluorescence imaging. Development 142, 4168–4179.

Lampropoulos, A., Sutikovic, Z., Wenzl, C., Maegele, I., Lohmann, J. U. and Forner, J. (2013). GreenGate---a novel, versatile, and efficient cloning system for plant transgenesis. PLoS One 8, e83043.

Landrein, B. and Hamant, O. (2013). How mechanical stress controls microtubule behavior and morphogenesis in plants: history, experiments and revisited theories. Plant J. 75, 324–338.

Lau, O. S. and Bergmann, D. C. (2012). Stomatal development: a plant’s perspective on cell polarity, cell fate transitions and intercellular communication. Development 139, 3683–3692.

Lee, L. R. and Bergmann, D. C. (2019). The plant stomatal lineage at a glance. J. Cell Sci. 132,.

Linder, H. P., Lehmann, C. E. R., Archibald, S., Osborne, C. P. and Richardson, D. M. (2018). Global grass (Poaceae) success underpinned by traits facilitating colonization, persistence and habitat transformation. Biol. Rev. Camb. Philos. Soc. 93, 1125– 1144.

Liu, T., Ohashi-Ito, K. and Bergmann, D. C. (2009). Orthologs of Arabidopsis thaliana stomatal bHLH genes and regulation of stomatal development in grasses. Development 136, 2265–2276.

Liu, L., Ashraf, M. A., Morrow, T. and Facette, M. (2024). Sto-Dual role of BdMUTE during stomatal development in the model grass Brachypodium distachyon matal closure in maize is mediated by subsidiary cells and the PAN2 receptor. New Phytol. 241, 1130–1143.

Lucas, W. J., Bouché-Pillon, S., Jackson, D. P., Nguyen, L., Baker, L., Ding, B. and Hake, S. (1995). Selective trafficking of KNOTTED1 homeodomain protein and its mRNA through plasmodesmata. Science 270, 1980–1983.

Lupanga, U., Röhrich, R., Askani, J., Hilmer, S., Kiefer, C., Krebs, M., Kanazawa, T., Ueda, T. and Schumacher, K. (2020). The Arabidopsis V-ATPase is localized to the TGN/EE via a seed plant-specific motif. Elife 9, e60568.

MacAlister, C. A. and Bergmann, D. C. (2011). Sequence and function of basic helix-loop-helix proteins required for stomatal development in Arabidopsis are deeply conserved in land plants. Evol. Dev. 13, 182–192.

MacAlister, C. A., Ohashi-Ito, K. and Bergmann, D. C. (2007). Transcription factor control of asymmetric cell divisions that establish the stomatal lineage. Nature 445, 537–540.

Matos, J. L., Lau, O. S., Hachez, C., Cruz-Ramírez, A., Scheres, B. and Bergmann, D. C. (2014). Irreversible fate commitment in the Arabidopsis stomatal lineage requires a FAMA and RETINOBLASTOMA-RELATED module. Elife 3, e03271.

McKown, K. H. and Bergmann, D. C. (2020). Stomatal development in the grasses: lessons from models and crops (and crop models). New Phytol. 227, 1636–1648.

McKown, K. H., Anleu Gil, M. X., Mair, A., Xu, S.-L., Raissig, M. T. and Bergmann, D. C. (2023). Expanded roles and divergent regulation of FAMA in Brachypodium and Arabidopsis stomatal development. Plant Cell 35, 756–775.

Mills, B. R. (2022). MetBrewer: Color Palettes Inspired by Works at the Metropolitan Museum of Art.

Nguyen, T.-H. and Blatt, M. R. (2024). Surrounded by luxury: The necessities of subsidiary cells. Plant Cell Environ.

Nunes, T. D. G., Zhang, D. and Raissig, M. T. (2020). Form, development and function of grass stomata. Plant J. 101, 780–799.

Nunes, T. D. G., Slawinska, M. W., Lindner, H. and Raissig, T. (2022). Quantitative effects of environmental variation on stomatal anatomy and gas exchange in a grass model. Quantitative Plant Biology 3, e6.

Nunes, T. D. G., Berg, L. S., Slawinska, M. W., Zhang, D., Redt, L., Sibout, R., Vogel, J. P., Laudencia-Chingcuanco, D., Jesenofsky, B., Lindner, H., et al. (2023). Regulation of hair cell and stomatal size by a hair cell-specific peroxidase in the grass Brachypodium distachyon. Curr. Biol. 33, 1844–1854.e6.

Pillitteri, L. J., Sloan, D. B., Bogenschutz, N. L. and Torii, K. U. (2007). Termination of asymmetric cell division and differentiation of stomata. Nature 445, 501–505.

Raissig, M. T. and Woods, D. P. (2022). Chapter Two - The wild grass Brachypodium distachyon as a developmental model system. In Current Topics in Developmental Biology (ed. Goldstein, B.) and Srivastava, M.), pp. 33–71. Academic Press.

Raissig, M. T., Abrash, E., Bettadapur, A., Vogel, J. P. and Bergmann, D. C. (2016). Grasses use an alternatively wired bHLH transcription factor network to establish stomatal identity. Proc. Natl. Acad. Sci. U. S. A. 113, 8326–8331.

Raissig, M. T., Matos, J. L., Anleu Gil, M. X., Kornfeld, A., Bettadapur, A., Abrash, E., Allison, H. R., Badgley, G., Vogel, J. P., Berry, J. A., et al. (2017). Mobile MUTE specifies subsidiary cells to build physiologically improved grass stomata. Science 355, 1215–1218.

Raschke, K. and Fellows, M. P. (1971). Stomatal movement in Zea mays: Shuttle of potassium and chloride between guard cells and subsidiary cells. Planta 101, 296–316.

R Core Team and R Foundation for Statistical Computing, Vienna, Austria (2023). R: A language and environment for statistical computing.

Renzaglia, K. S., Browning, W. B. and Merced, A. (2020). With Over 60 Independent Losses, Stomata Are Expendable in Mosses. Front. Plant Sci. 11, 567.

RStudio Team and RStudio, PBC, Boston, MA (2023). RStudio: Integrated Development Environment for R.

Rudall, P. J., Hilton, J. and Bateman, R. M. (2013). Several developmental and morphogenetic factors govern the evolution of stomatal patterning in land plants. New Phytol. 200, 598–614.

Rudall, P. J., Chen, E. D. and Cullen, E. (2017). Evolution and development of monocot stomata. Am. J. Bot.

Rui, Y., Chen, Y., Yi, H., Purzycki, T., Puri, V. M. and Anderson, C. T. (2019). Synergistic Pectin Degradation and Guard Cell Pressurization Underlie Stomatal Pore Formation. Plant Physiol. 180, 66–77.

Sack, F. D. (1994). Structure of the Stomatal Complex of the Monocot Flagellaria indica. Am. J. Bot. 81, 339–344.

Schindelin, J., Arganda-Carreras, I., Frise, E., Kaynig, V., Longair, M., Pietzsch, T., Preibisch, S., Rueden, C., Saalfeld, S., Schmid, B., et al. (2012). Fiji: an open-source platform for biological-image analysis. Nat. Methods 9, 676–682.

Serna, L. (2020). The Role of Grass MUTE Orthologues During Stomatal Development. Front. Plant Sci. 11, 55.

Serna, L. (2021). The Role of Grass MUTE Orthologs in GMC Progression and GC Morphogenesis. Front. Plant Sci. 12, 944.

Sharma, N. (2017). Leaf clearing protocol to observe stomata and other cells on leaf surface. Bio Protoc. 7, e2538.

Sozzani, R., Cui, H., Moreno-Risueno, M. A., Busch, W., Van Norman, J. M., Vernoux, T., Brady, S. M., Dewitte, W., Murray, J. A. H. and Benfey, P. N. (2010). Spatiotemporal regulation of cell-cycle genes by SHORTROOT links patterning and growth. Nature 466, 128–132.

Spiegelhalder, R. P. and Raissig, M. T. (2021). Morphology made for movement: formation of diverse stomatal guard cells. Curr. Opin. Plant Biol. 63, 102090.

Stebbins, G. L. and Shah, S. S. (1960). Developmental Studies of Cell Differentiation in the Epidermis of Monocotyledons. II. Cytological Features of Stomatal Development in the Gramineae. Dev. Biol. 2, 477–500.

Ursache, R., Andersen, T. G., Marhavý, P. and Geldner, N. (2018). A protocol for combining fluorescent proteins with histological stains for diverse cell wall components. Plant J. 93, 399– 412.

Wang, H., Guo, S., Qiao, X., Guo, J., Li, Z., Zhou, Y., Bai, S., Gao, Z., Wang, D., Wang, P., et al. (2019). BZU2/Zm-MUTE controls symmetrical division of guard mother cell and specifies neighbor cell fate in maize. PLoS Genet. 15, e1008377.

Weimer, A. K., Matos, J. L., Sharma, N., Patell, F., Murray, J. A. H., Dewitte, W. and Bergmann, D. C. (2018). Lineage- and stage-specific expressed CYCD7;1 coordinates the single symmetric division that creates stomatal guard cells. Development 145, dev160671.

Wickham, H. and Bryan, J. (2023). readxl: Read Excel Files.

Wickham, H., Averick, M., Bryan, J., Chang, W., McGowan, L., François, R., Grolemund, G., Hayes, A., Henry, L., Hester, J., et al. (2019). Welcome to the tidyverse. J. Open Source Softw. 4, 1686.

Wilke, C. O. and Wiernik, B. M. (2022). ggtext: Improved Text Rendering Support for “ggplot2.”

Wu, Z., Chen, L., Yu, Q., Zhou, W., Gou, X., Li, J. and Hou, S. (2019). Multiple transcriptional factors control stomata development in rice. New Phytol. 223, 220–232.

Yi, H., Chen, Y., Wang, J. Z., Puri, V. M. and Anderson, C. T. (2019). The stomatal flexoskeleton: how the biomechanics of guard cell walls animate an elastic pressure vessel. J. Exp. Bot. 70, 3561–3572.

Zhang, X., Facette, M., Humphries, J. A., Shen, Z., Park, Y., Sutimantanapi, D., Sylvester, A. W., Briggs, S. P. and Smith, L. G. (2012). Identification of PAN2 by Quantitative Proteomics as a Leucine-Rich Repeat–Receptor-Like Kinase Acting Upstream of PAN1 to Polarize Cell Division in Maize. Plant Cell 24, 4577–4589.

Zhang, D., Spiegelhalder, R. P., Abrash, E. B., Nunes, T. D. G., Hidalgo, I., Anleu Gil, M. X., Jesenofsky, B., Lindner, H., Bergmann, D. C. and Raissig, M. T. (2022). Opposite polarity programs regulate asymmetric subsidiary cell divisions in grasses. eLife 11, e79913.

Zuch, D. T., Herrmann, A., Kim, E.-D. and Torii, K. U. (2023). Cell cycle dynamics during stomatal development: Window of MUTE action and ramification of its loss-of-function on an uncommitted precursor. Plant Cell Physiol.

