## Supplemental Figures S1 - S10 for "Dual role of BdMUTE during stomatal development in the model grass *Brachypodium distachyon*"

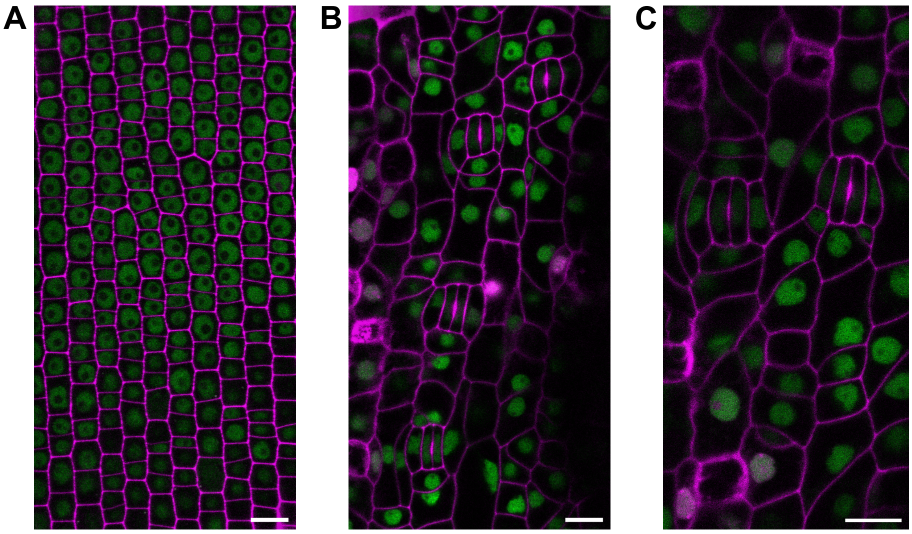


**Figure S1:** Ubiquitously expressed 3xGFP-BdMUTE in wild type (WT) background (WT*;ZmUBIp:3xGFP-MUTE*) induces ectopic divisions. (**A)** Early developmental zone of the leaf epidermis shows expression of 3xGFP-MUTE (green) in all cells. **(B, C)** Later stages of epidermal development show ectopic, subsidiary cell-like divisions. Shown are midplane confocal images of developmental zones in T0 lines stained with propidium iodide (PI, magenta). Scale bars = 10 $\mu$m.

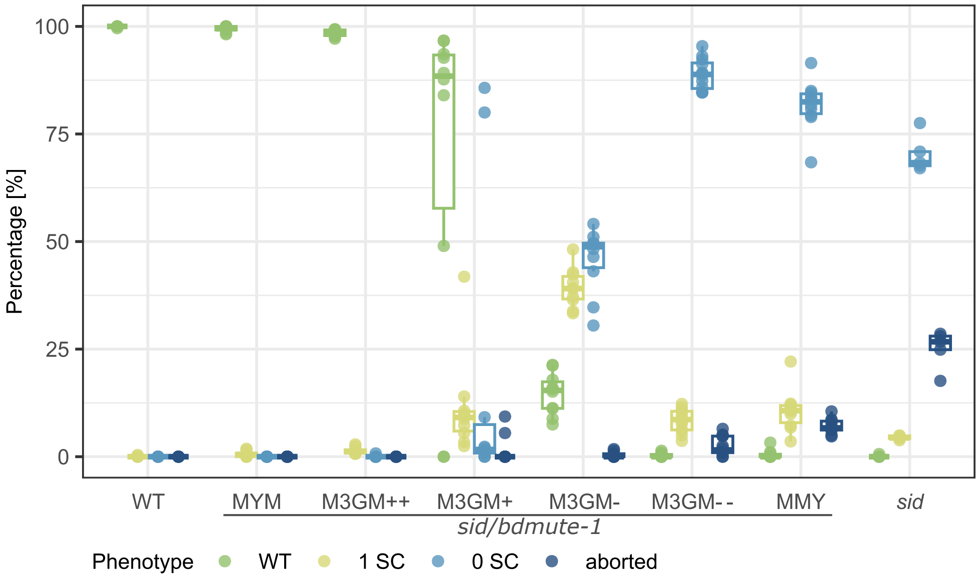


**Figure S2:** Complementation of *sid/bdmute-1* stomatal phenotypes by different complementation lines (same data as in Fig. 2E). Plant lines are wild type (WT), *sid/bdmute-1*;*BdMUTEp:YFP-BdMUTE* (MYM), four different *sid/bdmute-1*;*BdMUTEp:3xGFP-BdMUTE* lines (M3GM++, M3GM+, M3GM- and M3GM- -), *sid/bdmute-1*;*BdMUTEp:BdMUTE-YFP* (MMY) and *sid/bdmute-1*. Data from fully expanded $3^{rd}$ leaves of soil-grown plants 19-21 days after germination. Boxplots of data shown in Fig. 2E. n = 5-12 individuals per genotype and 864-1450 stomata per genotype. Each dot represents one individual.

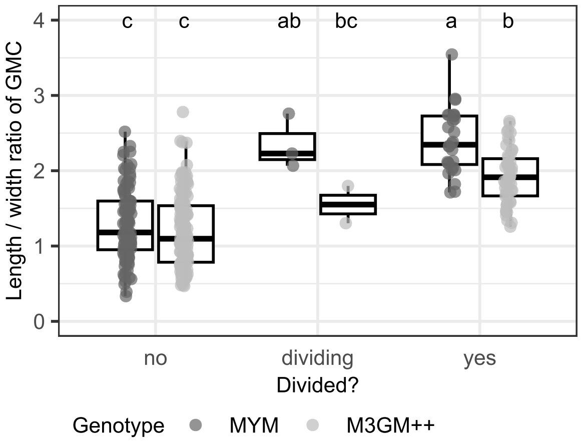


**Figure S3:** Length/width ratios of *sid/bdmute-1*;*BdMUTEp:YFP-BdMUTE* (MYM) and *sid/bdmute-1*;*BdMUTEp:3xGFP-BdMUTE* (M3GM++) guard cell complexes analyzed in Fig. 3E before, during and after guard mother cell (GMC) division. n = 2-6 individuals per genotype and 129-191 stomata per genotype. Each dot represents one stomatal complex. Significant differences are indicated with differing letters. Statistical test: ANOVA followed by Tukey's HSD test (alpha = 0.05).

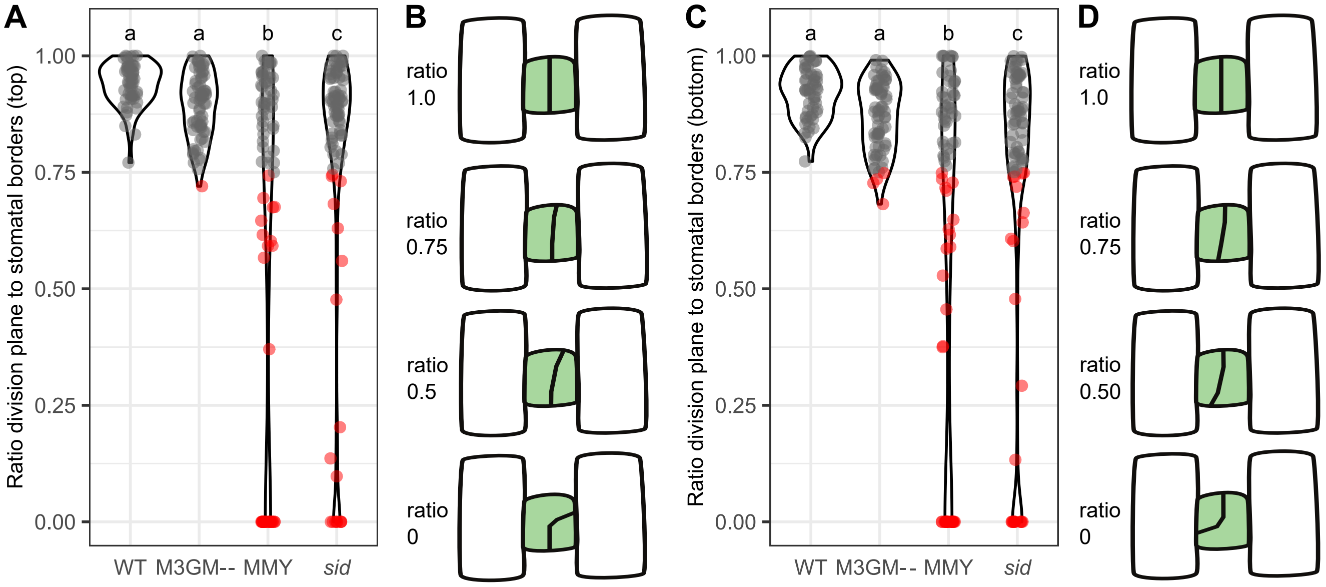


**Figure S4:** "Skewness" ratios of division plane orientation symmetry at the top and bottom of the guard mother cells (GMCs) in wild type (WT), *sid/bdmute-1*;*BdMUTEp:3xGFP-BdMUTE* (M3GM- -), *sid/bdmute-1*;*BdMUTEp:BdMUTE-YFP* (MMY) and *sid/bdmute-1*. Combined "skewness" ratios per stomatal complex are depicted in Fig. 4C. A straight line was drawn to connect the outward corners of the GMC at the top and bottom respectively (yellow line in Fig. 4B) and the distance from each corner to the intersection with the GMC division plane was measured along the previously drawn line (pink lines in Fig. 4B). The smaller distance was divided by the longer distance to obtain the ratio of division plane to stomatal borders plotted in **(A)** for the apical wall of the GMC. A ratio of 1 means the GMC divided longitudinally along the central plane, a ratio below 1 indicates a more skewed division orientation and at a ratio of 0 the division plane did not cross the upper stomatal border (i. e. transversal division). Gray dots indicate a ratio > 0.75 and red dots a ratio < 0.75. **(B)** Schematic representation of symmetric or skewed division plane orientation at the apical wall of the GMC with the respective ratios of division plane to stomatal borders. **(C)** Ratio of division plane to stomatal borders at the basal wall of the GMC. A ratio of 1 means the GMC divided longitudinally along the central plane, a ratio below 1 indicates a more skewed division orientation and at a ratio of 0 the division plane did not cross the lower stomatal border (i. e. transversal division). Gray dots indicate a ratio > 0.75 and red dots a ratio < 0.75. **(D)** Schematic representation of symmetric or skewed division plane orientation at the basal wall of the GMC with the respective ratios of division plane to stomatal borders. n = 7-8 individuals per genotype and 66-82 stomata per genotype (dots are stomata). Significant differences are indicated with differing letters. Statistical test: ANOVA followed by Tukey's HSD test (alpha = 0.05).

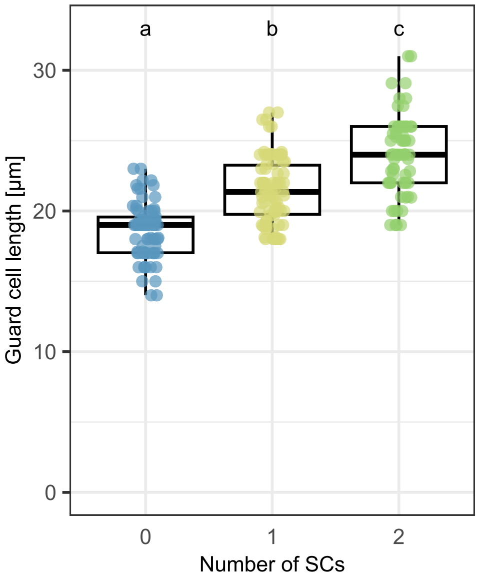


**Figure S5:** Guard cell length in *sid/bdmute-1*;*BdMUTEp:3xGFP-BdMUTE* (M3GM-) complexes with zero, one or two subsidiary cells (SCs). Same cells as measured in Fig. 5B. n = 2 individuals and 16-36 stomata per phenotype. Each dot represents one stomatal complex. Significant differences are indicated with differing letters. Statistical test: ANOVA followed by Tukey's HSD test (alpha = 0.05).

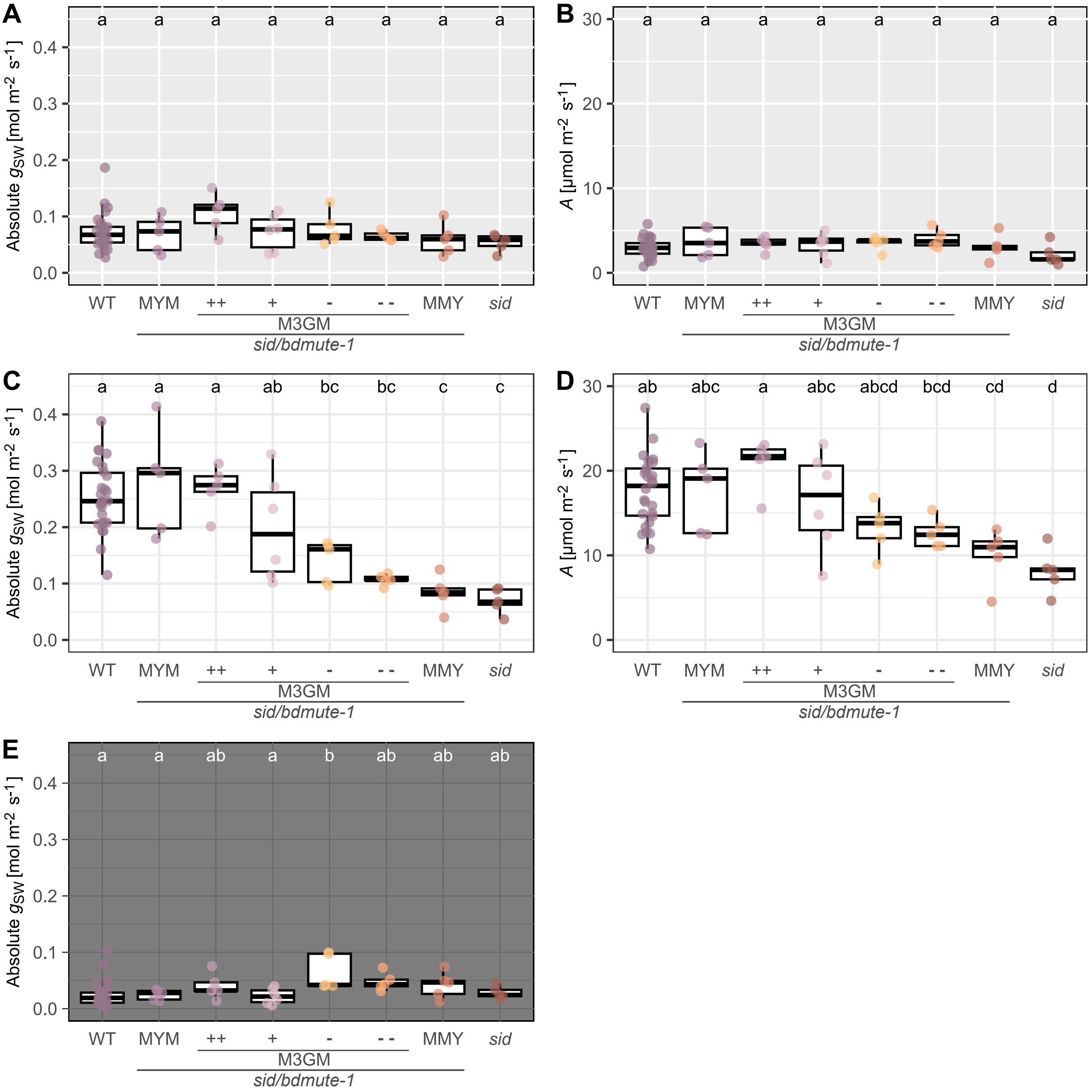


**Figure S6:** Steady state gas exchange values of measurements shown in Fig. 6 for wild type (WT), *sid/bdmute-1*;*BdMUTEp:YFP-BdMUTE* (MYM), the four different *sid/bdmute-1*;*BdMUTEp:3xGFP-BdMUTE* (M3GM++ to - -) lines, *sid/bdmute-1*;*BdMUTEp:BdMUTE-YFP* (MMY) and *sid/bdmute-1*. Average value of 5 minutes at the end of each light intensity step. Corresponding light intensity step is indicated by background color: Light gray (100 μmol m$^{-2}$ s$^{-1}$), white (1000 μmol m$^{-2}$ s$^{-1}$), and dark gray (0 μmol m$^{-2}$ s$^{-1}$). **(A) (C) (E)** Absolute stomatal conductance ($g_{sw}$). **(B) (D)** Carbon assimilation (*A*). n = 5-6 individuals per genotype, 26 individuals for WT. Dots represent individuals. Significant differences are indicated with differing letters. Statistical test: ANOVA followed by Tukey's HSD test (alpha = 0.05).

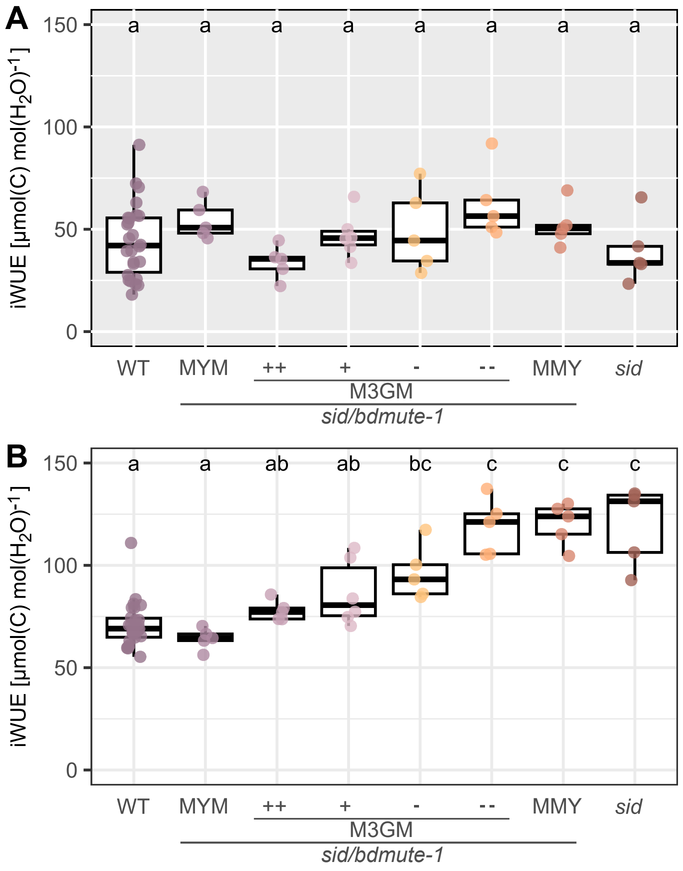


**Figure S7:** Steady state intrinsic water-use efficiency (iWUE) of measurements shown in Fig. 6 for wild type (WT), *sid/bdmute-1*;*BdMUTEp:YFP-Bd-MUTE* (MYM), the four different *sid/bdmute-1*;*BdMUTEp:3xGFP-BdMUTE* (M3GM++ to - -) lines, *sid/bdmute-1*;*BdMUTEp:BdMUTE-YFP* (MMY) and *sid/bdmute-1*. Average value of 5 minutes at the end of each light intensity step. Corresponding light intensity step is indicated by background color: **(A)** Light gray (100 $\mu$mol m$^{-2}$ s$^{-1}$), **(B)** white (1000 $\mu$mol m$^{-2}$ s$^{-1}$). n = 5-6 individuals per genotype, 26 individuals for WT. Dots represent individuals. Significant differences are indicated with differing letters. Statistical test: ANOVA followed by Tukey's HSD test (alpha = 0.05).

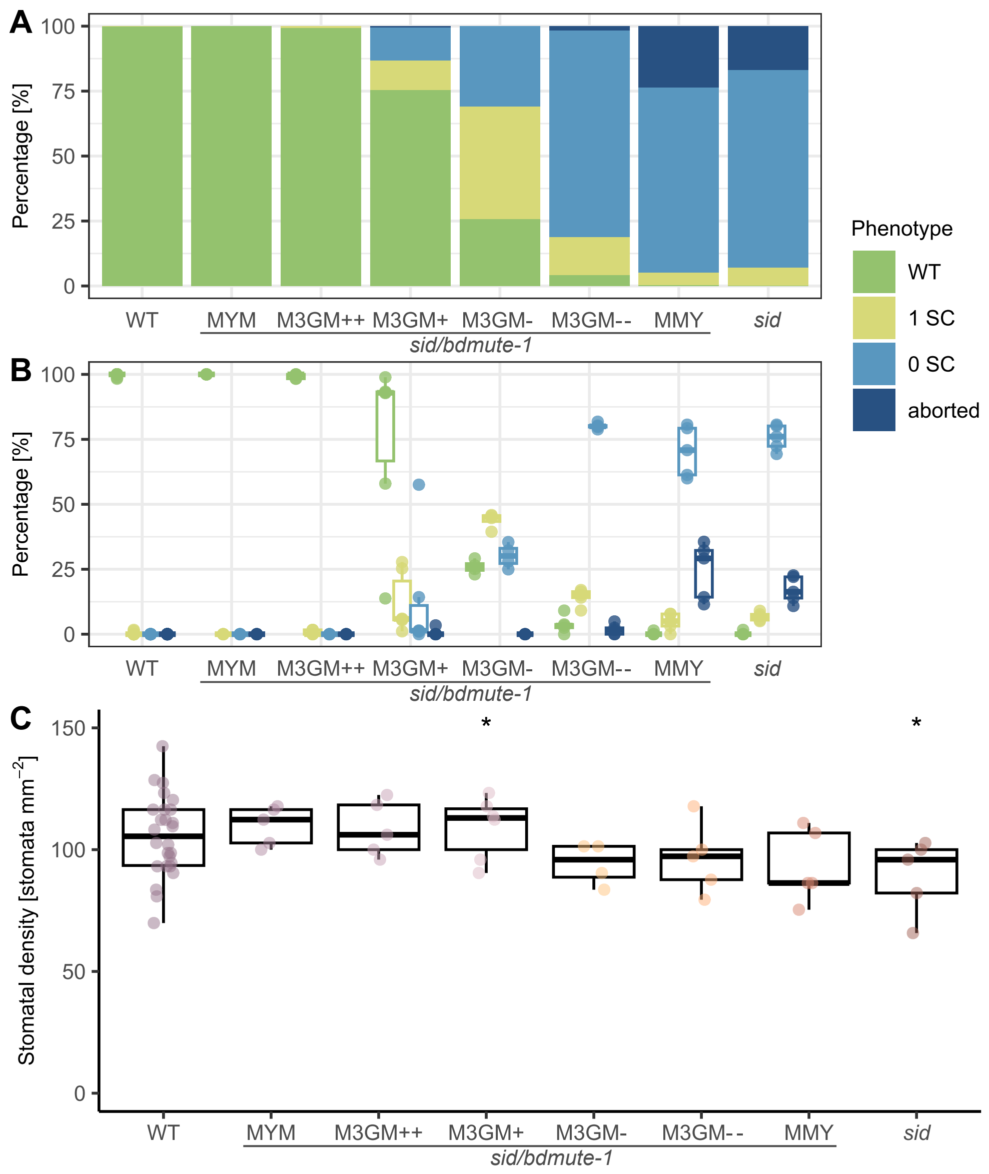


**Figure S8:** Stomatal morphology complementation phenotypes and stomatal density of leaves used for the leaf-level gas exchange measurements (Fig. 6) of wild type (WT), *sid/bdmute-1*;*BdMUTEp:YFP-BdMUTE* (MYM), the four different *sid/bdmute-1*;*BdMUTEp:3xGFP-BdMUTE* (M3GM++ to - -) lines, *sid/bdmute-1*;*BdMUTEp:BdMUTE-YFP* (MMY) and *sid/bdmute-1*. **(A)** Stacked bar plot of complementation of the *sid/bdmute-1* background in adult leaf. **(B)** Same data as in (A) depicted as box- and dot plot with dots representing individuals; n = 4-6 individuals per genotype, 26 individuals for WT. **(C)** Stomatal densities of the different lines; dots represent individuals. Asterisk indicates a significant difference to the WT individuals measured within the same experiment. n = 4-6 individuals per genotype, 26 individuals for WT, and 266-502 stomata per genotype. Statistical test: Two-sided Student's t-test (significant if $p < 0.05$) of the respective genotype compared to the WT individuals (n = 4-6) grown at the same time.

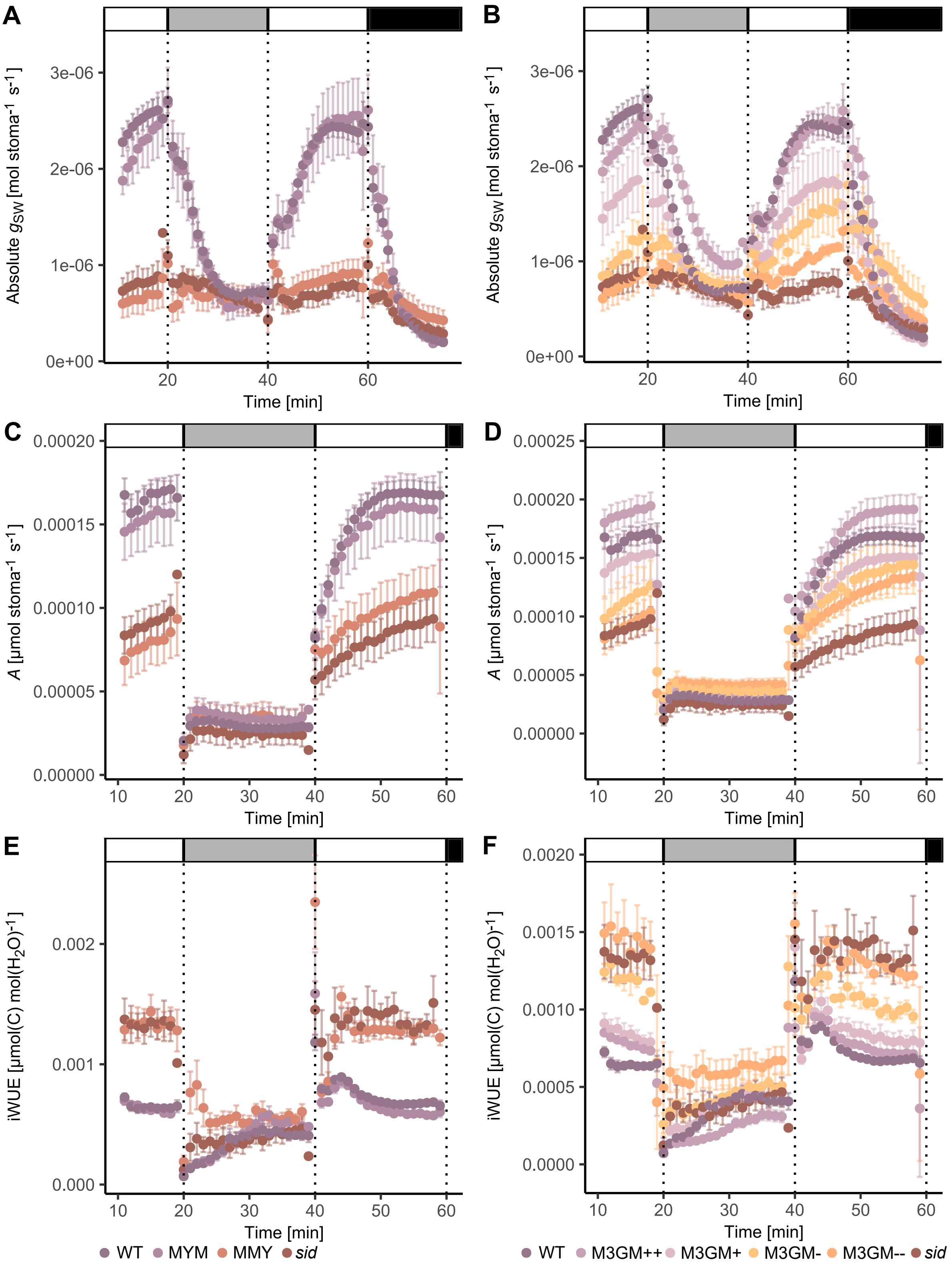


**Figure S9:** Leaf-level gas exchange values per stoma under changing light conditions. Data is the same as shown in Fig. 6 corrected by mean stomatal density per genotype. Changing light conditions are indicated by grayscale bars above each plot: White = 1000 $\mu$mol m$^{-2}$ s$^{-1}$, light gray = 100 $\mu$mol m$^{-2}$ s$^{-1}$, black = 0 $\mu$mol m$^{-2}$ s$^{-1}$. (**A**) Absolute stomatal conductance ($g_{sw}$) of wild type (WT), *sid/bdmute-1;BdMUTEp:YFP-BdMUTE* (MYM), *sid/bdmute-1;BdMUTEp:BdMUTE-YFP* (MMY) and *sid/bdmute-1*. (**B**) $g_{sw}$ of WT, the four different *sid/bdmute-1;BdMUTEp:3xGFP-BdMUTE* (M3GM) lines (++, +, - and - -) and *sid/bdmute-1*. (**C**) Carbon assimilation (*A*) of WT, MYM, MMY and *sid/bdmute-1*. (**D**) *A* of WT, M3GM lines and *sid/bdmute-1*. (**E**) Intrinsic water-use efficiency (iWUE) of WT, MYM, MMY and *sid/bdmute-1*. (**F**) iWUE of WT, M3GM lines and *sid/bdmute-1*. Note that the same WT and *sid/bdmute-1* data is shown in A, C, E and B, D, F. Measured were the youngest, fully expanded leaves of 3-4 week old, soil-grown plants; n = 5-6 individuals per genotype, 26 individuals for WT. Dots are means averaged across the individuals with error bars indicating standard error.

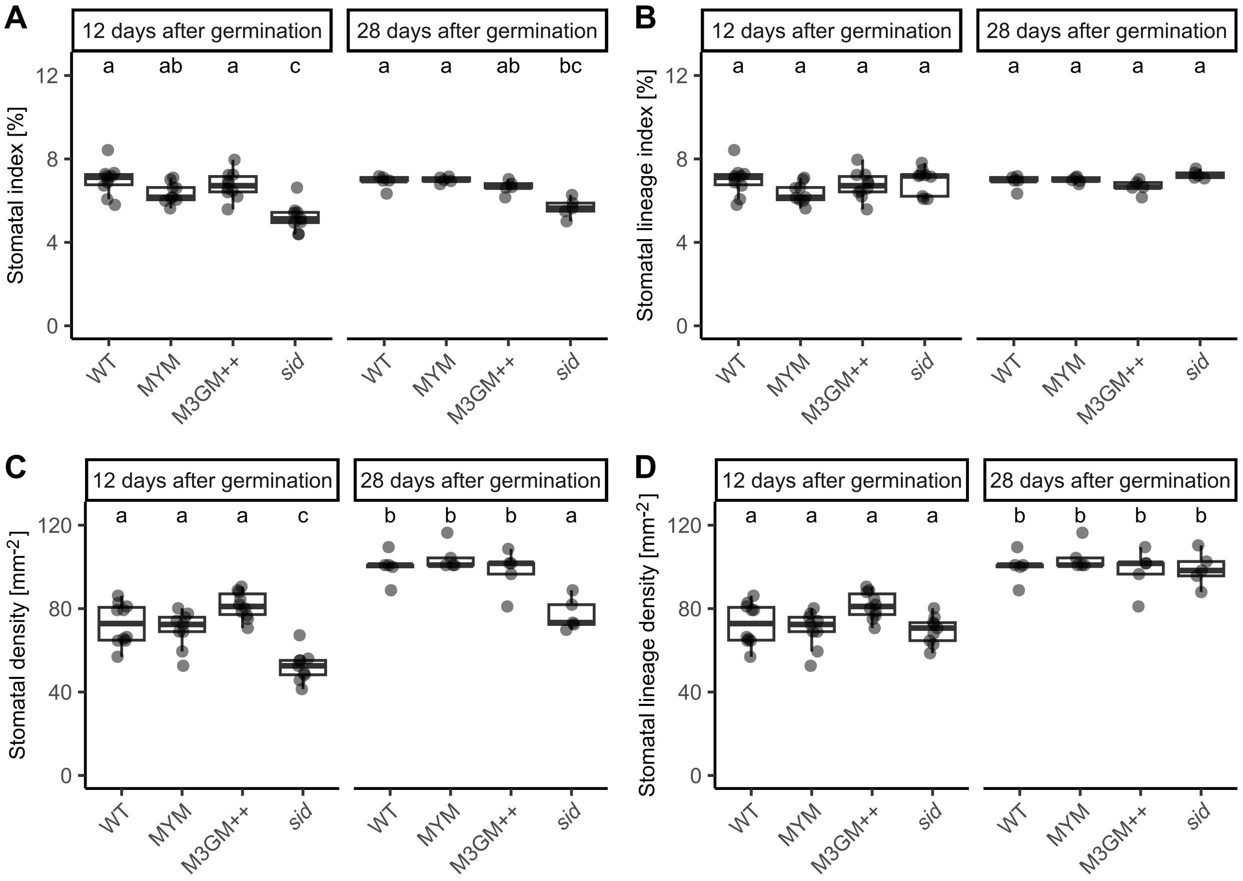


**Figure S10:** Stomatal index and stomatal density in wild type (WT), *sid/bdmute-1*;*BdMUTEp:YFP-BdMUTE* (MYM), *sid/bdmute-1*;*BdMUTEp:3xG-FP-BdMUTE* (M3GM++) and *sid/bdmute-1* leaves harvested 12 or 28 days after germination. **(A)** Stomatal index (number of stomata per total cell number). **(B)** Stomatal lineage index; this includes aborted stomatal complexes. **(C)** Stomatal density and **(D)** stomatal lineage density including aborted stomatal complexes. n = 9-10 individuals and 10'453-14'025 cells per genotype for 12 days after germination, 5 individuals and 7'906-8'651 cells per genotype for 28 days after germination. Each dot represents one individual.
